# Integrating SHAPE Probing with Direct RNA Nanopore Sequencing Reveals Dynamic RNA Structural Landscapes

**DOI:** 10.64898/2025.12.14.694260

**Authors:** J. White Bear, Grégoire De Bisschop, Éric Lécuyer, Jérôme Waldispühl

## Abstract

Traditional SHAPE experiments rely on averaged reactivities, which may limit information on folding patterns, alternate structures, and RNA dynamics. Short-read sequencing often suffers from false stopping, stalls, and biases during reverse transcription. The introduction of direct, long-read nanopore technology offers an opportunity to expand RNA structure probing methods to better understand RNA structural diversity. While many comparative approaches have been developed for detection of endogenous modifications, fewer have explored the expansion of SHAPE based methods. We introduce Dashing Turtle (DT), an algorithm using probabilistic, weighted, stacked ensemble learning to perform high-resolution detection of structural modifications that can capture detailed information about RNA architecture across dynamic structural landscapes. We apply our method to several well-characterized RNA samples, identify dominant conformations, and structurally conserved regions. We show that our landscapes correlate well with expected structures and recapitulate important functional elements. DT achieves accuracy 10–20% higher than comparable methods on many sequences. It accurately identifies structural features at a rate of 80–100%, approximately 10–30% better than its peers. DT’s predictions are robust across replicates and sub-sampled datasets and can help detect changes in conformational states, inform RNA folding mechanisms, and indicate interaction efficiency. Overall, it expands the capabilities of direct RNA sequencing and structural probing.

## Introduction

Current investigations of RNA structure often rely on the understanding of a single, final, low energy or native conformation to characterize the structure. However, RNA are highly flexible structures with a diversity of intermediate states and alternate folding hierarchies that exist to support interactions and other pathways in the free-energy landscape [1]. Indeed, ncRNAs have been shown to undergo multi-step conformational transitions that serve diverse functions including facilitating interactions with other molecules [2]. Alternative conformations, which may be higher energy, are often less explored in computational and experimental studies and characterization of dynamic states at thermodynamic equilibrium is difficult as most experimental methods require static or isolated molecules. Yet, it is understood that RNA enter alternative conformations as a transitional state, and can become kinetically trapped even in the presence of facilitating ions. These states exist in fractional populations, and are difficult and expensive to capture with approaches like cryogenic electron microscopy (cryo-EM), X-ray crystallagrophy, and nuclear magnetic resonance spectroscopy (NMR) [3, 1, 4, 5, 2].

Selective 2^′^-Hydroxyl Acylation analyzed by Primer Extension (SHAPE) based approaches are widely used experimental methods for probing secondary structure using reverse transcription (RT). SHAPE uses chemical reagents to bind to specific nucleotides or the 2^′^-hydroxyl ribose of the RNA backbone when it is most accessible in either unpaired positions or geometrically accessible base paired positions. Base-paired nucleotides usually represent stronger, hydrogen bonds between nucleotides that can make the 2^′^-hydroxyl ribose inaccessible. The chemical reagents form adducts using strong covalent bonds formed by acylation, a primer is added for DNA synthesis and the RNA is reverse transcribed to produce a complementary DNA or cDNA. The RNA adducts cause stops in reverse-transcription or mutations in the cDNA. Reverse-transcription stops can be analyzed by Capillary Electrophoresis (SHAPE-CE). Alternatively, next generation sequencing allows to profile cDNA mutations (SHAPE-MaP) [6]. Both methods use an averaged mutational profile over scDNA segments and can suffer from premature stopping of transcription, upstream and downstream effects, and can be difficult to MaP to varying isoform transcripts which decreases the structural resolution and obscures structural variation. Moreover endogenous modifications and structural features are lost or greatly obscured when transcribed [7, 6, 8]. Structural information about the heterogeneity of wider RNA dynamics are still challenging to obtain from current experimental approaches and is primarily approximated using ensemble or Boltzmann based energy landscapes [3, 1, 4, 5, 9]. Next generation sequencing platforms (NGS), such as Illumina, using cDNA and RT can introduce biases in cDNA synthesis by favoring regions of RNA that are less structured and more accessible, potentially underrepresenting regions with complex secondary structures. Secondary structures such as stem loops can also impede the enzyme’s progression, leading to incomplete cDNA synthesis or mis-priming [10].

The introduction of Oxford Nanopore Technology (ONT) offers the opportunity to expand the available mechanisms for RNA structure probing with chemical reagents using direct RNA sequencing. It offers improved isoform mapping and direct detection of endogenous and exogenous modifications [11, 12, 13, 14]. However, the ONT platform is inherently noisy and has a 98.7% basecaller accuracy rate at sequencing quality scores above 19 which has improved with the release of Dorado [15]. Comparatively, Illumina’s 99% accuracy at the same score, which increases to 99.9% at quality score of 30 [16]. These factors can make direct detection of RNA modifications quite challenging. Initial work by Aw et al. was able to show correlation between SHAPE based reactivity and ONT reactivity using common reagents like 2-(Azidomethyl)nicotinic acid imidazolide (NAI-N3), and 1-Methyl-7-nitroisatoic anhydride (1M7). However, these reagents significantly reduce the number and quality of sequenced reads and, the signal perturbations such as increased translocation time are not strongly predictive of modifications due to the high signal to noise ratio. Indeed, observed signal perturbations often showed high reactivity in bases that were not chemically modified and exhibited both upstream and downstream effects [11]. Stephensen et al. presented an alternate approach using Acetylimadizole (AcIm), a significantly smaller reagent, to modify and detect nucleotide bases using translocation time, or the amount of time each k-mer, or short sequence of nucleotides, resided in the nanopore. Again, correlation between increased translocation time and chemical modification was established, but the overall predictive power of the method was hindered by a low signal to noise ratio. Increased translocation times could also observed to be associated with certain bases even without modification, eg guanine [17]. Translocation times can also be effected by the compaction of the RNA structure prior to entering the nanopore or hybridization of the single strand after passing through the nanopore [18, 19, 20]. Newer methods rely primarily on the variance in electrical signal rather than translocation time. Neither method demonstrated single molecule nucleotide predictions or attempted to deconvolute the underlying RNA dynamics and focused on averaged mutational profiles [11, 17].

More recent methods attempt to apply Tombo, a package supported by ONT, that leverages probabilistic analysis of signal analysis to detect modifications using log likelihood ratio under a normal assumption, which has been useful for endogenous modifications, like *m*^6^a, but struggles on exogenous chemical modifications. Chemical adducts can induce distributions and random signal perturbations that are difficult to characterize under normal or Gaussian assumptions (Section 2.8.3). Adducts can cause heavy tailed distributions, asymmetry, multi-modality (multiple peaks, e.g. from alternative structures or stochastic dwell times), and are nonstationary (variance or mean drifts over time) causing bias, high false positives, and poor calibration across read batches or pores. Indeed, neither signal distributions (Appendix 6) nor positional distributions (Appendix 7) are represented by a normal Gaussian in the sequences we examined [21, 22, 12]. Current methods have shown modifications to adenines and cytosines using either dimethyl sulfate (DMS) or diethyl pyrocarbonate (DEPC) [22, 12]. Nano-Dms-MaP uses cDNA, not direct RNA, to detect mutations with DMS on adenine and cytosine bases, but, as we have discussed, cDNA can introduce significant biases, requires additional processing, and reverse transcription must be mediated by specialized cDNA to provide long transcripts of long reads [10, 12]. cDNA has preferential binding, so structural heterogeneity is easily lost [8, 23, 24]. Despite the use of cDNA, the Bohn et al. still note that noise is a significant issue in the signal and their method requires strict quality scores thresholds for each position in each read, ignoring indel (insertion and deletion) errors, and focuses only on mismatches in the DNA basecalling [12]. Bizuayehu et al. show a direct RNA sequencing approach, SMS-seq, using DEPC and Tombo to detect modifications of adenine bases. Adjusted mutual information is, then, used to infer long range dependencies of single molecules. This approach requires specific concentrations of DEPC to be selected with a trade off in read depth and synthetic RNA with precisely annotated adenine bases to establish thresholds and as we have already stated Tombo statistical approach does not generalize well to non-Gaussian distributions and assumes endogenous modifications with minimal interference with single nucleotide distributions [22, 21]. These recent works underline the significant effect of combined noise and relatively lower accuracy. Indeed, many recent work emphasizes the strong need for specialized algorithms to improve detection [22, 12, 11, 17].

To address these challenges, we introduce Dashing Turtle (DT) a probabilistic, stacked, ensemble learning method that incorporates multiple domains of analysis to capture structural modifications in nanopore data. DT is a novel specialized algorithm that produces results commensurate with traditional SHAPE based methods and extends to all four nucleotide bases. It’s detection is probabilistic and contextual, mirroring the contextual nature of biological data. DT offers high resolution single molecule detection that enables the investigation of the underlying heterogeneity in RNA structures. We apply our approach on well-characterized *in vitro* RNA transcripts and *in vivo* transcripts to demonstrate that DT reproduces and detects known structural variation of the transcripts in the structural landscape. We are able to identify dominant conformations and investigate alternative conformations that may be obscured by averaged mutational profiling.

## Material & Methods

### In vitro transcription

DNA templates were PCR-amplified from plasmid or minigene using Q5 DNA polymerase with the primers listed in Appendix Table 8. Templates for T. thermophila riboswitch, HCV IRES, E. coli tmRNA and HSPA1A were amplified from plasmid pT7L-21, dl HCV 1b, pJW28-tmRNA, and MV04-HSP70, respectively. Templates for lysine riboswitch and FMN riboswitch were amplified from minigenes indicated in Appendix Table 6. Template for hc16 ligase was generated by PCR using the Q5 polymerase using tiled oligonucleotides (Appendix Table 8) following the procedure described in [25]. Reverse primers for SHAPE-CE experiments additionally contained a 3’ cassette as described in [24] to allow reverse transcription primer binding. RNAs were in vitro transcribed using T7 RNA polymerase following the manufacturer’s instructions for 2 h at 37°C. Template DNA was digested with DNAse I for 15 minutes at 37°C and the resulting RNAs were purified using RNAClean XP beads. RNA integrity was assessed by gel electrophoresis and RNA concentration was measured using 260nm absorbance. RNA for direct sequencing were further polyadenylated using a poly(A) polymerase for 20 minutes at 37°C. Poly(A) RNA were purified using RNAClean XP beads and tail addition was confirmed by gel electrophoresis.

### Chemical Probing

RNA was folded in 40 *µ*L of folding buffer (60 mM HEPES, pH 7; 140 mM NaCl; 10 mM KCl) by denaturation for 1 minute at 95^°^C, snap-cooling on ice, and incubation for 30 minutes at 37^°^C (or 42^°^C for the heat-shock experiment) after addition of 1 mM MgCl_2_. Samples were then split into two aliquots. One aliquot was treated with 0.75 M AcIm to reach a final concentration of 75 mM, while the other was treated with the same volume of DMSO as the non-modified control. DMS probing was performed similarly, using a final concentration of 100 mM DMS and neat ethanol for the non-modified control. Samples were incubated for 10 minutes at 37^°^C (or 42^°^C for the heat-shock experiment) for modification.

For SHAPE-MaP, an additional denaturing control was obtained by denaturing RNA at 95^°^C in denaturing buffer (25% formamide, 50 mM HEPES, pH 8, and 4 mM EDTA) and modifying for 1 minute at 95^°^C after addition of 75 mM AcIm. Samples were subsequently purified using RNAClean XP beads. Elution was performed in 10 *µ*L H_2_O, and RNA concentration was measured using a Qubit fluorimeter. Modified RNA was then analyzed by SHAPE-CE, SHAPE-MaP, or direct RNA sequencing as described in the following sections, along with their unmodified controls.

### SHAPE-MaP

SHAPE-MaP was performed as described in using the small RNA workflow [6]. The protocol was applied to RNA from 3 individual probing experiments. RNA were reverse transcribed using the reverse primers from step 1 PCR primer pairs listed in Appendix Table 7. Reverse-transcription products were amplified by 5 cycles of PCR using Q5 DNA polymerase. The products were purified using PureLink PCR micro spin columns. Amplicons were further amplified by a 25 cycle barcoding PCR and purified using Agencourt AMPure XP beads. After measuring concentration on a Qubit fluorimeter and bioanalyzer quality control, amplicons were pooled and sent for sequencing on a MiSeq 500 Nano V2 chip at the IRIC facility. Triplicate data were processed using Shapemapper2 [26] using the –amplicon option and the parameter –min-depth set to 1000.

### SHAPE-CE

SHAPE-CE was performed as in [27]. In brief, RNA were reverse transcribed using the MMLV reverse transcriptase using either a 6-FAM-labeled primer or a NED-labeled primer and 0.5 mM ddTTP for the sequencing reaction. cDNA from both reactions were then mixed and precipitated with 0.3 M sodium acetate, 1 *µg* glycogen and 75% ethanol, resuspended in HiDi formamide and submitted to capillary electrophoresis. Fluorescence signals were processed using QuSHAPE [7] and peak areas were normalized using the IPANEMAP suite [28] (*https://github.com/Sargueil-CiTCoM/ipasuite*) to determine reactivities. Reactivities were averaged from 3 individual probing experiments.

### Direct RNA sequencing

Polyadenylated RNA were processed for ONT sequencing using SQK-RNA002 kit. 200 ng of RNA were ligated using the provided RTA adapter following the manufacturer instructions and reverse-transcribed using Supercript III reverse transcriptase. Ligated, reverse-transcribed RNA were loaded on Flongle flow cells (R9.4.1 chemistry) and run until the number of active pore dropped below 5.

### Reagents

Chemicals used in this study were 1-Acetylimidazole (Sigma-Aldrich, St. Louis, MO, USA; Cat. No. 157864-25G), dimethyl sulfide (Sigma-Aldrich, St. Louis, MO, USA; Cat. No. 75-18-3), and dimethyl sulfoxide (DMSO) (Sigma-Aldrich, St. Louis, MO, USA; Cat. No. D8418-100ML). Enzymes included T7 RNA polymerase (New England Biolabs, Ipswich, MA, USA; Cat. No. M0251L). Critical commercial assays and kits were the Direct RNA Sequencing Kit (Oxford Nanopore Technologies, Oxford, UK; Cat. No. SQK-RNA002) and Flongle flow cells (Oxford Nanopore Technologies, Oxford, UK; Cat. No. FLO-FLG001). Reagents are listed in Appendix 6

### Biological Resources

The full list of oligonucleotides is listed in Appendix Table 6 and Appendix Table 7.

### RNA Constructs

Table 1 lists all RNA constructs, organisms, the reference structures, and the number of transcripts obtained, both unmodified and modified, used in this analysis. For HCV IRES and *E. coli* tmRNA structures are referenced from described below. The HCV IRES construct corresponds to PDB entry 3T4B, encompassing nucleotides 116-324 of the 382 nucleotide RNA provided by Bruno Sargueil. This region includes a 21 nucleotide crystallization module that replaces the native stem loop spanning positions 132-276, as described by Berry *et al*. [29], Fig.1. The *E. coli* tmRNA construct corresponds to PDB entry 3IZ4 (chain 1), which matches the experimental RNA used in this study except for the replacement of loop nucleotides 118-126 with an MS2 hairpin tag introduced for sample preparation, as described by Fu *et al*. [30].

**Fig. 1.**
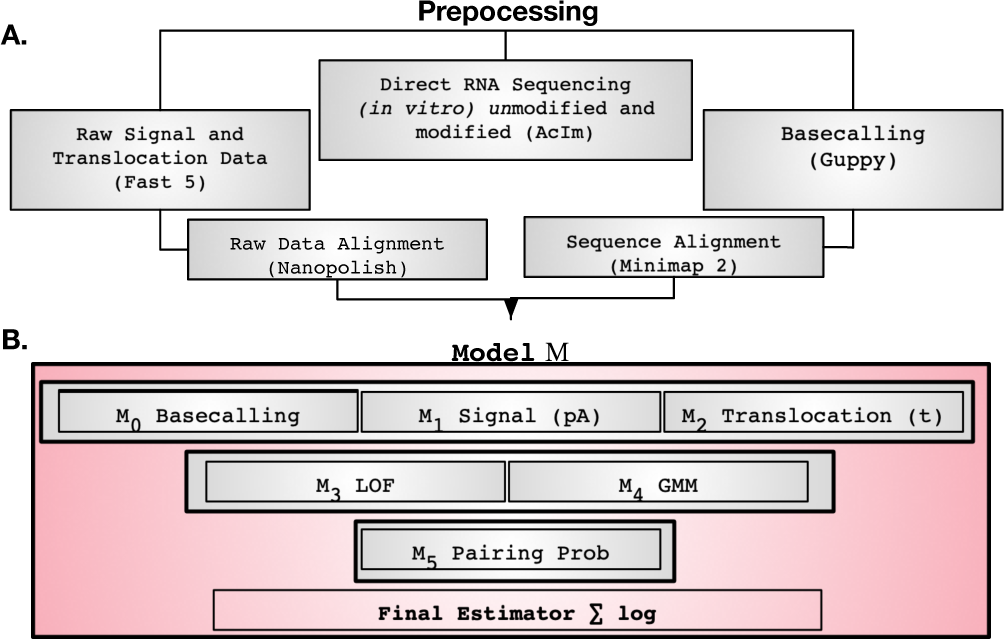
Figure shows the preprocessing steps for biological sequencing and data extraction from ONT pipeline. The aligned raw data include signal, translocation times, and basecalling data serve as the main inputs into the Model *M* (A.). The lower panel describes the ensemble Model *M* ‘s primary components *M*_0_, …, *M*_5_ used to predict base modifications for each read and position of an RNA sequence. *M*_0_, *M*_1_, *M*_2_ is a comparative analysis of peaks in the decomposed basecalling, signal, and translocation data analyzed using the LNPPD algorithm (Section 2.8.4). *M*_3_ and *M*_4_ describe machine learning components used to capture wider sequence context (Section 2.8.6, 2.8.7) and *M*_5_ which uses the base pairing probability calculated from the stacking energies computed using RNAFold [9]. The final estimator represents the calculation of the reactivity function described in Section 2.8.9 (B.)

**Table 1.** Summarized read counts per RNA sequence, including full species names, experimentally determined PDB structures where available, and number of both unmodified and modified transcripts obtained. Structures referenced from literature are cited.

| Sequence Name | Species | PDB Entry | Unmodified Reads | Modified Reads |
| --- | --- | --- | --- | --- |
| Bacterial Ribonuclease P RNA (RNase P) 16 | *Geobacillus stearothermophilus* | 2A64 | 261,617 | 74,406 |
| Class-hc ligase ribozyme (hc16 Ligase) RNA | Synthetic | 6WLO | 144,192 | 34,140 |
| E. coli tmRNA (ssrA) | *Escherichia coli* | [30] | 122,088 | 5,839 |
| Tetrahymena thermophila Group I Intron Ribozyme (L-21 ScaI variant) | *Tetrahymena thermophila* | 7EZ0 | 79,672 | 9,611 |
| HCV IRES RNA | *Hepacivirus C* | [29] | 35,077 | 8,999 |
| Lysine Riboswitch RNA | *Bacillus subtilis* | 3D0X | 1,044 | 1,277 |
| Flavin mononucleotide (FMN) Riboswitch RNA | *Bacillus subtilis* | 3F2Q | 189 | 309 |

### Experimental Design

We first sequenced RNA with ONT and basecalled with Guppy [15, 31]. Unmodified RNA transcripts were used as a comparative control. Next, we applied Acetylimidazole (AcIm) to transcripts to acylate bases and measure reactivity. AcIm, as identified by Stephenson et al., is a 2^′^-hydroxyl acylation analyzed by primer extension (SHAPE) which generates a small enough adduct to pass through nanopore after reacting with accessible structures of the ribose on the RNA backbone [17]. AcIm is not base specific and identifies any accessible area of the RNA molecule, so it can be used to probe all four nucleotides. It is also much smaller than traditional SHAPE probes to allow it to more easily pass through nanopores without significant blockage. While we did see a reduction in the number of reads from nanopore after modification, we where still able to obtain a statistically significant sample.

The exact sample size (n) for each experimental group/condition are listed in Table 1 for each sequence. Where the number of modified reads exceed 6,000, a random sample of 6,000 reads was used as the modified data to decrease computation time. Controls sequences were used for validation and compared to contemporary methods and experiments. This data was combined with several methods and a weighting scheme to modulate the noise and determine the likelihood that a nucleotide has an AcIm adduct.

In total our training set includes ≈ 800, 000 transcripts combined without chemical modification and ≈ 100, 000 transcripts modified with AcIm. Finally, we use the information about modification patterns of each molecule to generate a structural landscape that models the diversity of structural conformations in the control sample and performed a comparative analysis of structural and functional features. We, then, performed this analysis on our test transcript, and compared the results to existing literature (Table 1).

### Statistical Analyses

#### Stacked Ensemble Learning Model

Ensemble learning combines multiple base learners and has been shown to outperform individual algorithms in several cases. By aggregating the predictions of several models, ensemble methods can reduce variance, mitigate bias, and improve overall predictive performance across different metrics. Common ensemble strategies include bagging, which reduces variance by averaging multiple models trained on bootstrapped samples; boosting, which sequentially trains models to correct previous errors and reduce bias; and stacking, which learns an optimal combination of diverse base learners [32, 33, 34].

ONT platforms while very useful for sequence analysis exhibit several challenges to standard machine learning approaches: the data output are very noisy and non-stationary. ONT is a biological nanopore with variation for each sequence or read across each nanopore and variation in the amount of (picoAmperes) *pA* and translocation time, *t*, required to determine the base in the nanopore [15]. These issues result in significant variation in the data even for unmodified RNA sequences. Predicting structural modification in this context is, also, primarily an unsupervised task. An ablation study was performed to assess the best parameters for our model (Appendix A.1). Below we describe a novel ensemble learning model that combines signal processing and machine learning, denoted *M*, used to decompose the signals and predict the modifications needed to define the landscape.

#### Preprocessing

Signal data obtained from the Fast5 files generated by Guppy were aligned with current measurements per read and nucleotide position using Nanopolish [35, 31]. In our kit, there were approximately 1024 five-mers, representing all possible five nucleotide regions that could occupy a nanopore and a reference range of average signals associated with each five-mer [15]. These values are resolved only to the level of basecalling, the addition of a structure probe changes the expected average signal based on the underlying structure of the RNA (single, base paired, loop, kinks, twists), the nucleotide being modified (A,C,G,U), and the chemical reagent being used to probe the RNA. Additionally, the raw signal drifts over time in a non-stationary manner. Thus, raw signal data requires additional preprocessing step to normalize nanopore signals.

#### Addressing non-stationarity

ONT signals represent time series from individual nanopores, each of which produces independent current traces, and multiple nanopores are embedded in each membrane. These signals are known to drift over time [36, 37]. We observed that, when combined across reads using standard normalization, the mean and variance of nanopore signals were non-stationary processes [38], which we confirmed using the Augmented Dickey–Fuller (ADF) and Kwiatkowski–Phillips–Schmidt–Shin (KPSS) tests [39, 40]. A standard log transform of the signal features was insufficient to remove this non-stationarity.

To allow for comparison across non-stationary signal data, we robustly scaled each read. Let *S* = {*s*_1_, *s*_2_, …, *s*_*n*_} denote all reads for a given sequence, and let *X*_*i*_ represent the robustly scaled signal for read *s*_*i*_, such that *X*_*i*_ ∼ *N* (0, 1). We assumed independence between reads, as each nanopore sequences a single molecule at a time. By the Central Limit Theorem [41], the sum of independent random variables, even if not perfectly Gaussian, tends toward a normal distribution as *n* increases. Thus, given independent (approximately normal) random variables *X*_1_, …, *X*_*n*_ with means 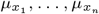 and variances 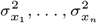, the sum

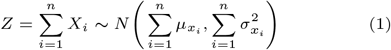

is approximately normally distributed. We used *Z* as a generalized normalization across reads for each sequence and confirmed stationarity of *Z* using the ADF and KPSS tests [39, 40].

#### Basecall and Signal Decomposition

##### *M*_0_: Detecting Modifications in Basecall Analysis

Basecalling determines the nucleotides A, C, G, or U in each sequence. Previous works have used errors in insertions, deletions, or mismatches in comparative analyses to detect endogenous modifications such as *m*6*A* [42, 43]. Here, we take comparative reactivities as a component of the ensemble model, *M*_0_.

For a given RNA sequence *S* and all reads *s* ∈ *S*, let *s*_*i*_ denote a single read containing all bases at positions 1, …, *n*, where *n* is the sequence length. Let *k* be the number of times a given position is read; note that each position in a single read can be read multiple times. The basecall reactivity for read *i* is defined as a vector of length *n*, with each element representing the sum of insertions *I*, deletions *D*, and mismatches *M* at that position, divided by the number of reads covering that position:

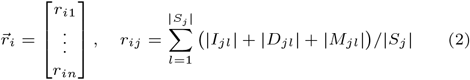

We define the basecall reactivity difference for read *i* as

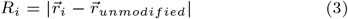

where 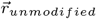 is the average reactivity across all unmodified reads.

#### Signal and Translocation Reactivity

Signal analysis represents the average signal (ionic current, *pA*) for the central base in the five-mer occupying the nanopore. The translocation time represents the dwell time, i.e., the duration a base remains in the nanopore for a given five-mer.

We define a reactivity vector 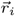 for read *i*, where *x* is either the signal (*pA*) for *M*_1_ or dwell time *t* for *M*_2_, *i* indexes reads, and *j* indexes positions along the sequence with |*j*| being the total number of positions. For each position *j*, repeated measurements of the signal or dwell time are averaged; the total number of repeated measurements at position *j* in read *i* is denoted *k*_*j*_. The first matrix represents the measurements from a single modified read, from which the mean over all unmodified reads is subtracted. The absolute value of this difference defines the reactivity per read, *r*_*i*_(*x*), as Equation 4:

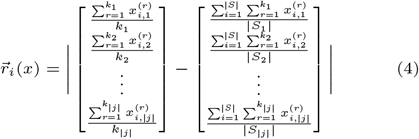

Here, *i* = 1, …, |*S*| indexes individual reads, and *j* = 1, …, |*j*| indexes positions along the sequence, where |*j*| is the total length of the sequence. The variable 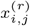 represents the *r*-th repeated measurement of the signal (in picoamperes) or the translocation time for read *i* at position *j*, and *k*_*j*_ is the total number of repeated measurements at position *j* in read *i*. The quantity |*S*_*j*_ | denotes the total number of reads covering position *j*. Thus, 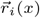 represents the position reactivity for each read and position, as the absolute deviation from the mean signal/translocation time of unmodified reads. The signal or translocation reactivity for read *i* in model *M* − 1 can then be defined similar to basecall reactivity as Equation 5:

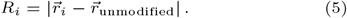

For the specific case of translocation time *t* in model *M*_2_, we apply a weighting factor *λ* to account for the fact that guanine (G) bases typically exhibit longer dwell times than other nucleotides. This was shown by Stephensen et al. and confirmed in signal and translocation time of our control sequences [17]. Thus, if the central base is a G, the probability *P* (*M*_2_) is reduced by *λ* which was scaled to the level of bias introduced in signal and translocation times by G in such a way that G did not trigger false positive modifications in unmodified data. If it is not a G, *P* (*M*_2_) is increased. However, if *M*_2_ does not exhibit an increase in dwell time, we decrease the overall weight *ω* of the joint probability across *M*_0_–*M*_2_ (Equation 14). Empirically, while increased translocation time was not definitive evidence of chemical modification, the absence of increased translocation time was strongly indicative of an unmodified site. The weighting parameters *λ* and *ω* were then optimized using control (unmodified) sequences.

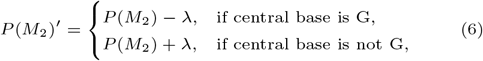

##### Analysis

This comparative analysis incorporates background subtraction to remove some of the noise from the signal and allows differences in signal and translocation time in modified sequences to be more clearly represented. After background subtraction, we apply peak detection using previously developed algorithm: localized, non-parametric probabilistic detection (LNPPD [44]). LNPPD addresses limitations of noisy non-Gaussian distributions in nanopore data using Gaussian kernel density estimate (KDE) to estimate local relative probabilities (within 3-5 nucleotide) for each base. LNPPD demonstrates improved accuracy of between 12% and 22% of detected modified bases across all sequences compared to other methods. While significant basecall errors associated with a position or where LNPPD indicates a high probability of modification are useful for distinguishing noise from true modification, leveraging the decomposition into three independent signals is important improves detection compared to single model approaches Independently, each of these signals can show significant variation due to the stochasticity of nanopore basecalling, but the joint probability, *ω*[*P* (*M*_0_) * *P* (*M*_1_) * *λP* (*M*_2_)], is less likely to be the result of noise. The caveat for these models is that they are highly localized, which could represent a local drift in the signal rather than modification. LNPPD does not take into account the context of the wider sequence. We examine the wider context by continuing with a stacked, ensemble learning model that offers better long range contextualization and generalize across sequences [44].

#### Cross Read Distributions

For wider contextual information, we evaluated control sequences using several unsupervised/ semi supervised methods on modified RNA to detect modifications and selected the algorithms with the best performance. We applied Local Outlier Factor (LOF) using novelty detection to determine modified bases in the decomposed signal analysis data for both *pA* and *t* defined in Section 2.8.5. While *M*_0_-*M*_2_ can detect comparative differences in very short 2-3 base ranges using LNPPD, LOF employs density based estimation using local neighborhoods of feature vectors derived from each nucleotide’s signal and translocation characteristics. Each nucleotide in each read is represented as a point in feature space, and neighborhoods are formed from the *k* nearest points, which can include nucleotides from different reads and different positions with similar signal and translocation patterns. This allows LOF to capture population level anomalies, flag rare modifications or unusual translocation events, and expand the localization beyond the 2-3 base range, removing data that is atypical locally but typical in the context of the neighborhood [45]. For *M*_3_, we implemented the Local Outlier Factor (LOF) algorithm using the scikit-learn framework [46] which computes a local density-based anomaly score for each nucleotide’s feature vector. Each nucleotide *j* in read *r*_*i*_ was represented as a feature vector 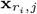 comprising the current and translocation time, *pA* and *t* respectively.The *k* nearest neighbors are identified to form the local neighborhood 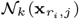. The local reachability density (LRD) is defined as the inverse of the average reachability distance to these neighbors:

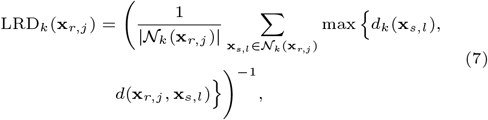

where *d*_*k*_(**x**_*s,l*_) is the distance from neighbor **x**_*s,l*_ to its *k*th nearest neighbor, and *d*(**x**_*r,j*_, **x**_*s,l*_) is the Euclidean distance between feature vectors. The LOF score for nucleotide *j* in read *r* is then:

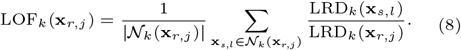

In scikit-learn, the negative LOF score is used for novelty detection, with more negative values indicating stronger deviation from the local neighborhood and possible modification. For each read *r*, the LOF predictions across all nucleotide positions are collected into vectors 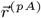 and 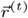 for the two signal components. The probability of modification for each read is computed element-wise:

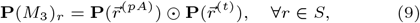

where ⊙ denotes element-wise multiplication and 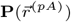 and 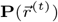 are vectors of probabilities derived from the LOF predictions for each nucleotide in read *r*. This approach enables detection of rare signal or translocation anomalies across multiple reads and positions, extending the localization beyond short-range 2–3 base comparisons captured by models *M*_0_–*M*_2_. LOF performed at 50-60% area under the curve (AUC) and offered increased accuracy in detection of outliers in the data for neighborhoods across multiple reads outside of a positional context.

#### Positional Distributions and Correlations

For *M*_4_, we employed a Gaussian Mixture Model (GMM) [47, 48] to model the joint distribution of *pA* and *t* at each nucleotide position *j* across all reads. Comparatively, LOF does not maintain the sequential boundaries, GMM strongly models these correlations. A GMM was trained on unmodified sequences, allowing the model to capture correlations between *pA* and *t* rather than treating them as independent decomposed variables. The joint distribution of *pA* and *t* at each nucleotide position *j* across all reads *r* ∈ *S*. Let **x**_*r,j*_ = (*pA*_*r,j*_, *t*_*r,j*_) denote the feature vector for nucleotide *j* in read *r*. Then, the GMM at position *j* is:

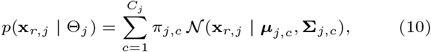

where:

- *C*_*j*_ is the number of Gaussian components at position *j*, optimized empirically;
- *π*_*j,c*_ are the mixture weights satisfying 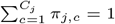;
- ***µ***_*j,c*_ and **Σ**_*j,c*_ are the mean vector and covariance matrix for component *c* at position *j*;
- 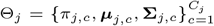 represents all GMM parameters for position *j*.

After training on unmodified reads, the probability of modification for nucleotide *j* in read *r* is computed as the complement of the likelihood under the learned GMM:

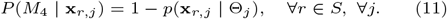

Vectorizing across all reads at position *j*, we can write:

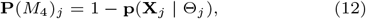

where **X**_*j*_ = *{***x**_*r,j*_ *}*_*r*∈*S*_ and **P**(*M*_4_)_*j*_ is the vector of modification probabilities for all reads at position *j*. These probabilities can then be combined with outputs from other models (*M*_0_–*M*_3_) in an ensemble framework.

In cases where modifications were strong and consistent across most reads, the resulting distributions could be well approximated by Gaussian components. Here, the GMM naturally transitioned from a unimodal distribution on unmodified sequences to clearly separated bi-modal clusters on modified sequences (Appendix 7). The number of Gaussian components was optimized empirically to maximize separation at positions known or predicted to be modified.

However, as we discussed, GMM performance decreases in scenarios where the signal distributions deviated substantially from Gaussian assumptions, such as when distributions were noisy or heavy-tailed (Appendix 6). In these cases, separation into distinct clusters did not occur (Appendix 7), and alternative models (*M*_0_–*M*_3_), which rely on non-parametric or density-based approaches, were more accurate. Indeed, after modeling the distributions at each position across all reads, we observed that a substantial fraction of positions exhibited non-Gaussian characteristics. While probabilistic bounds, such as those derived from Chebyshev’s inequality [49], could in principle generalize modification probabilities, non-parametric methods remained more reliable in practice. Across sequences, *M*_4_ achieved an area under the ROC curve (AUC) of approximately 60-70%, depending on sequence context. While it was one of the most accurate single algorithms on average, it also exhibited substantial variance in performance across different sequences.

#### Base Pairing Probabilities

To incorporate structural information into the ensemble reactivity model, we extended Equation 14 with a structure-aware correction term based on predicted pairing probabilities. Specifically, we define *P*_paired_ as the maximum probability of base pairing (or reduced reactivity) predicted by RNAFold [9]. For each read, base-pairing probabilities were computed for all nucleotides, considering both intra- and inter-molecular interactions, and calibrated using the predicted nanopore reactivities to refine the folding energy parameters. For each nucleotide, the bi-directional pairing probabilities (*P*_*i,j*_ and *P*_*j,i*_) were compared, and the maximum value was assigned to *P*_paired_.

The predicted ensemble reactivity for each base, *R*_pred_, was then compared against *P*_paired_ to determine the final reactivity, *R*_*j*_. When *R*_pred_ exceeded a high reactivity threshold, *τ*_*h*_ (typically 0.7 on a unit scale), the experimental signal was retained, ensuring that experimental reactivity dominates over predicted structural constraints. Conversely, when *R*_pred_ was below *τ*_*h*_ and the pairing probability exceeded a high-confidence threshold, *τ*_*p*_ (typically 0.97), the nucleotide was adjusted to a non-reactive state, reflecting strong base-pairing stability. This integration allows *P*_paired_ to modulate the ensemble reactivity term in Equation 14, balancing experimental evidence with predicted structural constraints. The adjustment rule is summarized in Equation 13.

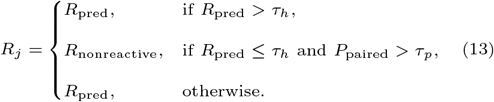

Typical threshold values are *τ*_*h*_ = 0.7 for high predicted reactivity and *τ*_*p*_ = 0.97 for strong pairing probability. This formulation ensures that experimentally strong reactivities override predicted pairing, while weak reactivities consistent with strong base pairing are suppressed to reflect structural stabilization. The resulting *R*_*j*_ thus represents a structure aware ensemble reactivity score that integrates both experimental and thermodynamic evidence.

#### Stacked Ensemble Layer

The stacked ensemble layer is the final estimator for the model, *M*, by optimizing a weight vector, 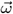, over all reads in the control set, and used the log-probability of the individual models to compute a final score representing the likelihood of nucleotide reactivity. We found that locally optimized weights 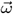 outperformed standard Bayesian or regression-based stacking approaches [32, 33, 34]. Let the reactivity, *R*, represent a vector of reactivities for each nucleotide position *j*. Equation 14 summarizes *P* (*M*_*k*_) representing the vectors of probabilities of modification for each nucleotide predicted by model *M*_*k*_:

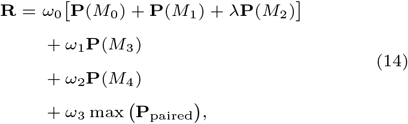

Here, *ω*_0_, *ω*_1_, *ω*_2_, *ω*_3_ are the optimized weights for each model or structural term, and *λ* allows for differential scaling of *P* (*M*_2_) (Section 2.8.5). The max(*P*_paired_) term ensures that positions with high predicted base-pairing probabilities contribute appropriately to the final reactivity score (Section 2.8.8). This weighted ensemble approach integrates both probabilistic model predictions and structural information, producing a robust and extensible estimator of RNA reactivity across all reads.

### Dashing Turtle Software

Dashing Turtle has been developed into a software package including the complete algorithm described in the text, predictions, and generation of landscapes described below. The package is available for Python 3.11 at and DashingTurtle on Anaconda. Full documentation for the software can be found at Dashing Turtle Installation Guide. Data preprocessing scripts are available at: Dashing Turtle Preprocess on GitHub. Source code for Dashing Turtle is available at: Dashing Turtle on GitHub.

## Results and Discussion

While average metrics on pooled transcripts can be less precise and do not reflect RNA dynamics, particularly in the case of nanopore based sequencing; we evaluated DT’s detection accuracy against comparative algorithms and experimental methods to measure the effectiveness of DT before constructing the landscape.

### Validation I: Probing Comparison

We probed five of our control RNA with AcIm and evaluated modifications using SHAPE-CE and SHAPE-MaP techniques. FMN and Lysine were not probed using SHAPE and reserved as a hold out for structural validation (Section 3.2). Next, we sequenced both probed and unprobed transcripts using nanopore and evaluated the modifications using DT. Then, we compared methods presented in Aw et al. using SVM based PORE-cupine [11], Stephensen et al.’s method using Kolmogrov-Smirnov based nanoSHAPE [17], and finally Vienna’s RNAcoFold using DT reactivities [9]. For RNAcoFold, we calculated the control molecule’s probability of pairing with itself, and another molecule in the sample and took the maximum base pairing scores across all ensembles for each nucleotide based on DT reactivities and applied a threshold of .97, where base pairing probabilities below this number were considered unmodified. We compared the predictions from each algorithm against the experimentally predicted modifications from both SHAPE-MaP and SHAPE-CE. We observed significant variation between the experiments. SHAPE-CE and SHAPE-MaP overlapped at a rate of about 72% (Figure 2 A.). To calculate the nanopore overlap we took the average agreement of all algorithms excluding RNAcoFold, with total agreement between all three methods observed at about 47%. Nanopore based methods predicted on average about 10% less. However, DT modifications were in agreement approximately 72% comparative to experimental method agreement (Table 9). DT’s predictions were also the most accurate overall compared to SHAPE-MaP and SHAPE-CE. Figure 2 D./E. show the accuracy of all algorithms predicting modifications on the five control sequences. Table 9) details the agreement between all respective methods, with DT overall having the highest percentage of bases in agreement. DT’s accuracy is the result of a conservative model and the predictions are less sensitive than comparative algorithms with very low fold change (Figure 2 B.). The detection threshold in DT is set for a probability of modification above .75 and actively excludes false positives or less reactive nucleotides. Comparative algorithms over predict modifications significantly almost doubling the rate of actual predictions (Figure 2). We compared the precision, and recall scores of the methods as well with all methods exhibiting high variance based on the sequence. Precision across all sequences reached a maximum of 67% (DT), 35% (SHAPE-CE), compared to SHAPE-MaP. Recall or sensitivity reached a maximum of 50% (nanoSHAPE) reflecting the high fold-change, and SHAPE-CE at 47% compared to SHAPE-MaP. In Figure 2 C. shows DT’s performance using an optimized threshold on *T. thermophila* across Accuracy 62%, Precision 21%, Sensitivity 39%, F1 27%, Pearson’s Correlation 5%, and Matthew’s Correlation Coefficient 5% (Appendix A.2). Modifications were consistent with regards to structural location (e.g. loop or bulge), but not to the extent of modification. Extent of modification can be described as the number bases modified within a structural location. For example, loop of size *k*, could contain up to *k* modifications. DT prioritizes accuracy so extent of modification, sensitivity, can be lower (Figure 2 C.). However, all methods saw significant variation in the extent of modification in a given structural location with the minimum being 1. It is difficult to determine if the extant is due to inaccessibility or a bias in the modification rate of the experiment. We measure the structural features annotated in our controls in the next section.

**Fig. 2.**
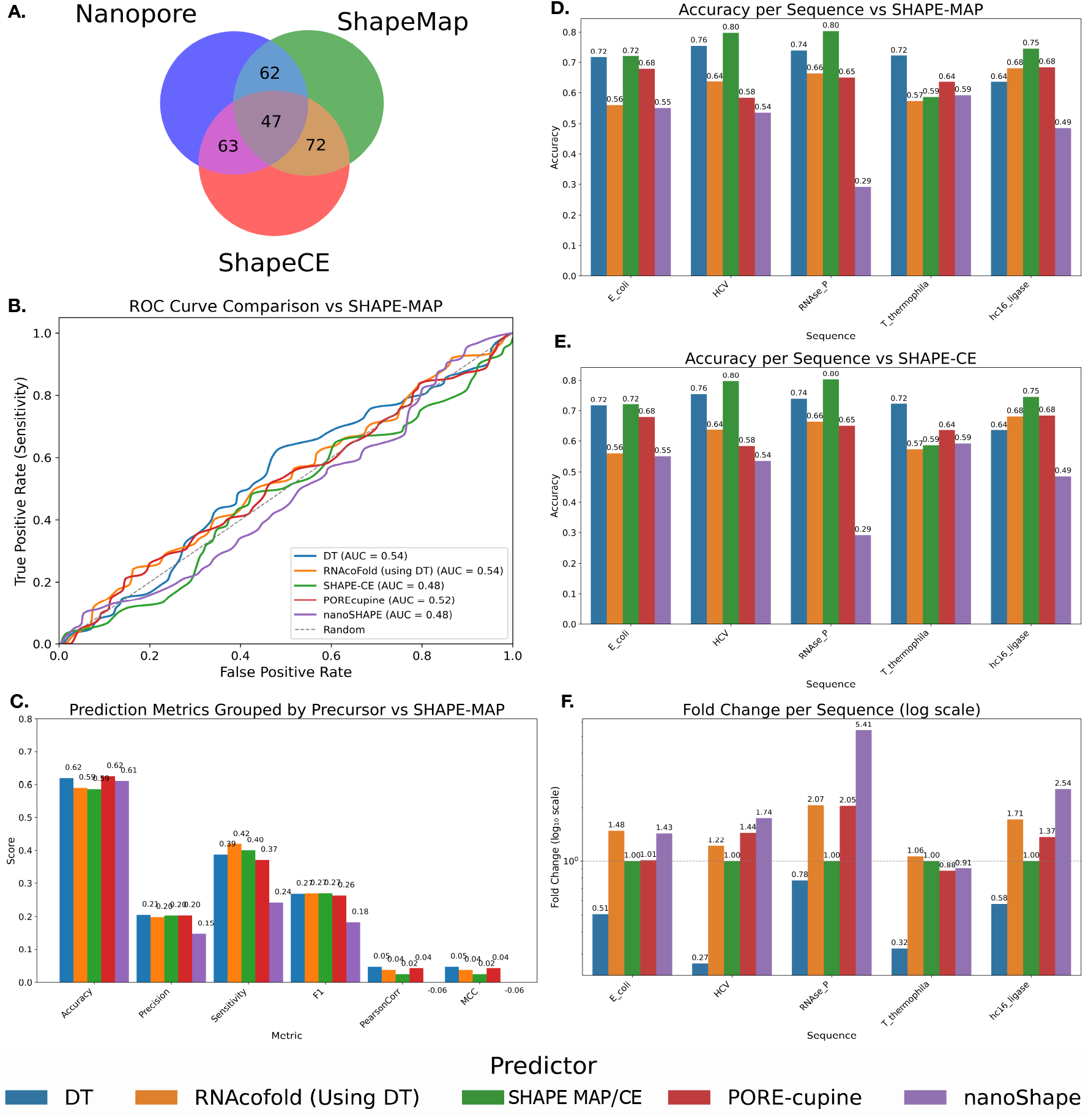
Comparison between SHAPE-MaP, SHAPE-CE, DT, PORE-cupine, and nanoSHAPE on control sequences *E. coli*, HCV, RNase P (*T. thermophila*), and hc16 ligase. (A) Venn diagram displaying the overlap between nucleotide modifications detected. SHAPE-MaP and SHAPE-CE agree on approximately 72% of detections. SHAPE-MaP and nanopore-based methods agree on approximately 62% of detections. SHAPE-CE and nanopore-based methods agree on approximately 63% of detections. All three methods agree on approximately 47% of detected modifications. Appendix Table 9 shows all percent overlaps and total predictions. (B) ROC Curve of each algorithm against SHAPE-MaP on *T. thermophila*. DT used an optimized threshold and performed at AUC 54% when predictions were compared against SHAPE-MaP, equivalent to RNAcoFold with DT reactivities, and has the highest AUC. SHAPE-CE had AUC of 48%, POREcupine 52 % and nanoSHAPE 48%. (C) Bar plot showing DT’s performance after doubling the Fold Change to extend coverage on *T. thermophila*. Accuracy against SHAPE-MaP decreased from 72% to 62%, Precision 21%, Sensitivity 39%, F1 27%, Pearson’s Correlation 5%, and Matthew’s Correlation Coefficient 5%. DT was still a top performer in all categories and tied for accuracy. *T. thermophila* is highly conserved functional structure so has fewer reactive positions across all metrics as observed in the lower experimental correlation between SHAPE-CE and SHAPE-MaP. (D) Bar plot showing the accuracy of each method compared to SHAPE-MaP. DT outperforms all other methods except SHAPE-CE for the hc16 ligase. (E) Bar plot showing the accuracy versus SHAPE-CE for each method on each control sequence. DT is the most accurate, excluding SHAPE-MaP for all sequences. (F) Bar plot showing the fold change (log scale) for each method on each sequence of the control. DT is noticeably conservative, while nanoSHAPE overpredicts in several instances by more than double the experimentally identified cases.

### Validation II: Structural Features

Based on experimentally confirmed structures, we next assessed performance by evaluating whether the predicted modified bases correlated with loops and bulges of the confirmed structures. Structural features are generally highly conserved in functional transcripts. However, the use of reverse transcription (RT) in traditional SHAPE based protocols is biased toward structurally optimal molecules which creates more uniform distributions for average modifications [50, 51, 52, 53, 54, 55, 56, 57]. ONT’s direct reads do not have a known structural bias in sequencing, meaning mis-folded structures may be sequenced at the same rate as energetically stable conformations. Extremely strong structures or very long molecules may still have reduced translocation efficiency through the pore, but will be sequenced at similar rates. Similarly, medium and long RNAs fold poorly without ions; native folding probability can drop to 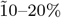 or less, especially for tertiary structures. This is reflected in the lower percentages of transcripts conforming to native structure (Section 3.2.2)and indicates that using average metrics on pooled transcripts is less precise and doesn’t accurately reflect RNA dynamics, particularly in the case of nanopore based sequencing [58, 59, 60]. Still we evaluated error rates of predicted modifications of base paired nucleotides, e.g. if a nucleotide is predicted to have high reactivity but is base paired in its native conformation then this is a Type 1 error or false positive. We calculated the total number of errors, removing base paired loops closures because AcIm is known to modify these base pairs at a higher rate and calculated the overall error [17]. On average SHAPE-CE exhibited the highest average error rates ≈ 9.04%, DT errors on average were ≈ 6.6%, and SHAPE-MaP error were ≈ 2.7% when compared to crystallized or NMR based structures [61]. SHAPE-MaP’s lower error rate could be attributed to two factors: ONT has a 98.7% accuracy rate at quality score above 19 [15] compared to Illumina’s 99% accuracy at the same score, which increases to 99.9% at quality score of 30 [16, 62, 63]. Table 2 shows each methods adherence to the known structures of our control sequences and measures the number of loops, and bulges detected. Our approach performed comparably to existing experimental methods, as suggested in Section 3.1. DT performed in the top category for detection in most instances and in all but one instance for bulge detection (Table 2 E). nanoShape while performing well has elevated fold changes and predicts most nucleotides as modified (Figure 2 F.); it also struggled comparatively with smaller features like bulges (Figure 3 B)). POREcupine offered a reasonable balance for this issue and performed second best overall. In general, the data suggests using nanopore based methods can increase detection of structural features significantly.

**Table 2.** Percent Accuracy with Structural Features. Loops and Bulges reference three metrics for all algorithms compared to the experimentally confirmed structure. Table displays the percent of modifications that agree with loops or bulges in the confirmed RNA structure by sequence. A loop requires at least two modified bases and includes the enclosing base pairs. A bulge requires at least one modified base in the enclosing nucleotides or the unpaired nucleotide. Probabilistic ensemble methods with nanopore perform significantly better across the board and annotate most major structural features, in some cases at double the rate of other methods. For bulge detections, it was the best performing method in all except one case. nanoSHAPE, while showing strong performance, relies on predicting most nucleotides as modified (see Fold Change Figure 2E) and has poorer separation and accuracy. The next best performers overall were POREcupine and RNAcoFold informed by DT reactivities. For FMN and Lysine, SHAPEMAP and SHAPECE were not performed, so the data is missing and this is indicated by the dash symbol. However, predictions from RNAcoFold, POREcupine, and nanoSHAPE were performed, and 0 indicates no detections.

| Sequence | DT |  | RNAcoFold<br>(DT reactivities) |  | POREcupine |  | nanoSHAPE |  | SHAPEMAP |  | SHAPECE |  |
| --- | --- | --- | --- | --- | --- | --- | --- | --- | --- | --- | --- | --- |
|  | Loop | Bulge | Loop | Bulge | Loop | Bulge | Loop | Bulge | Loop | Bulge | Loop | Bulge |
| *E. coli* | 85% | **83%** | 80% | 67% | 75% | 67% | **95%** | 67% | 65% | 50% | 80% | 50% |
| *T. thermophila* | **91%** | **75%** | 83% | 63% | 78% | 69% | 78% | 38% | 61% | 50% | 87% | 69% |
| HCV | 91% | **92%** | 73% | 54% | 77% | 69% | **95%** | 85% | 68% | 54% | 23% | 38% |
| RNAse P | 82% | 67% | 64% | 73% | 86% | 53% | **96%** | **93%** | 68% | 67% | 50% | 27% |
| hc16 ligase | **100%** | **100%** | 79% | 90% | 93% | 60% | 100% | 90% | 93% | 40% | 79% | 80% |
| FMN | **100%** | 0 | 83% | 0 | 83% | 0 | 100% | 0 | — | — | — | — |
| Lysine | **100%** | **50%** | 73% | 0 | 73% | 50% | 0 | 0 | — | — | — | — |

**Fig. 3.**
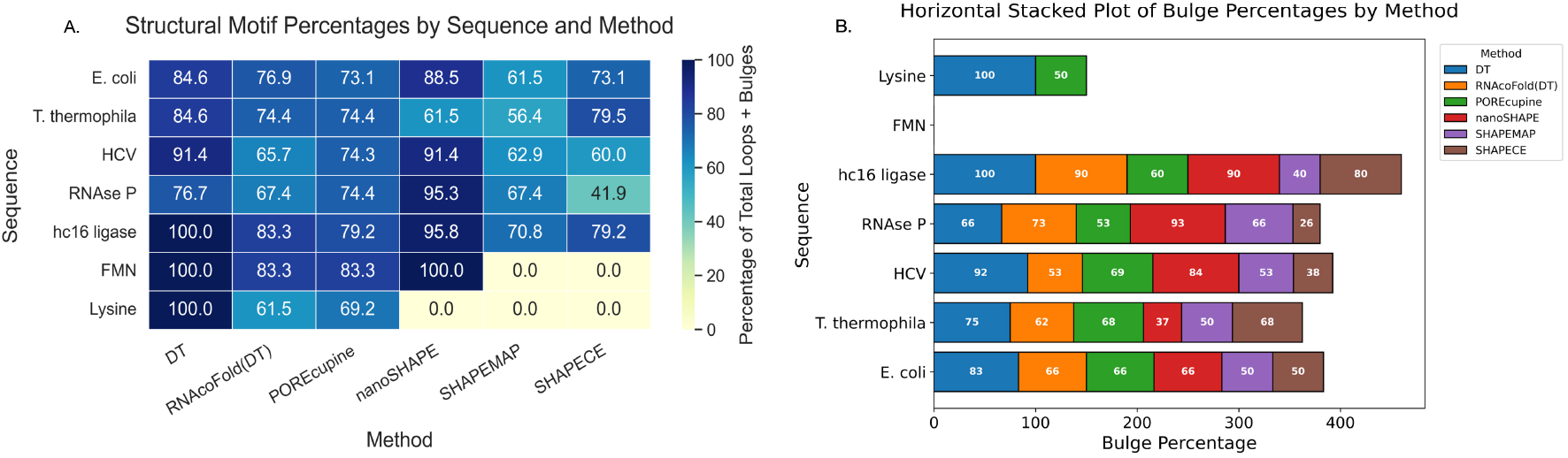
Comparison between SHAPE-MaP, SHAPE-CE, DT, PORE-cupine, and nanoSHAPE on control sequences *E. coli*, HCV, RNase P (*T. thermophila*), hc16 ligase, FMN, and Lysine. (A) Heatmap displays the aggregate percent of loops and bulges detection for each method. DT is able to detect most features and outperforms other methods. (B) A stacked bar plot of the bulges detected for each sequence for each method. DT is the top performer in all cases except one, RNAse P where nanoSHAPE performs better.

#### Generating the Landscape

After validating DT’s ability to accurately detect modifications and structural features, we generate the structural landscape which expose structural variations by obtaining the reactivity for each position and each read individually. Figure 4 A show’s the reactivity of reads for HCV IRES over the landscape (Table 1). Each landscape represents the reactivity of each *in vitro* transcripts modified with AcIm. The data suggests that highly accessible or functional positions important positions are more likely to remain so across the landscape, but significant variability can be observed. Variable reactivity implies that not all transcripts assume the expected native or low energy confirmation. The changes in modification could indicate random fluctuations; however, when this is repeated across multiple reads it becomes statistically significant. We can, then, view these patterns as possible alternative conformations in solution (Figure 5). Given that RNA are less likely to fold into native conformation when not in the presence of stabilizing ions and that RNA molecules can experience kinetic traps and fluctuations known as breathing, the data could be elucidating these patterns [1, 64].

**Fig. 4.**
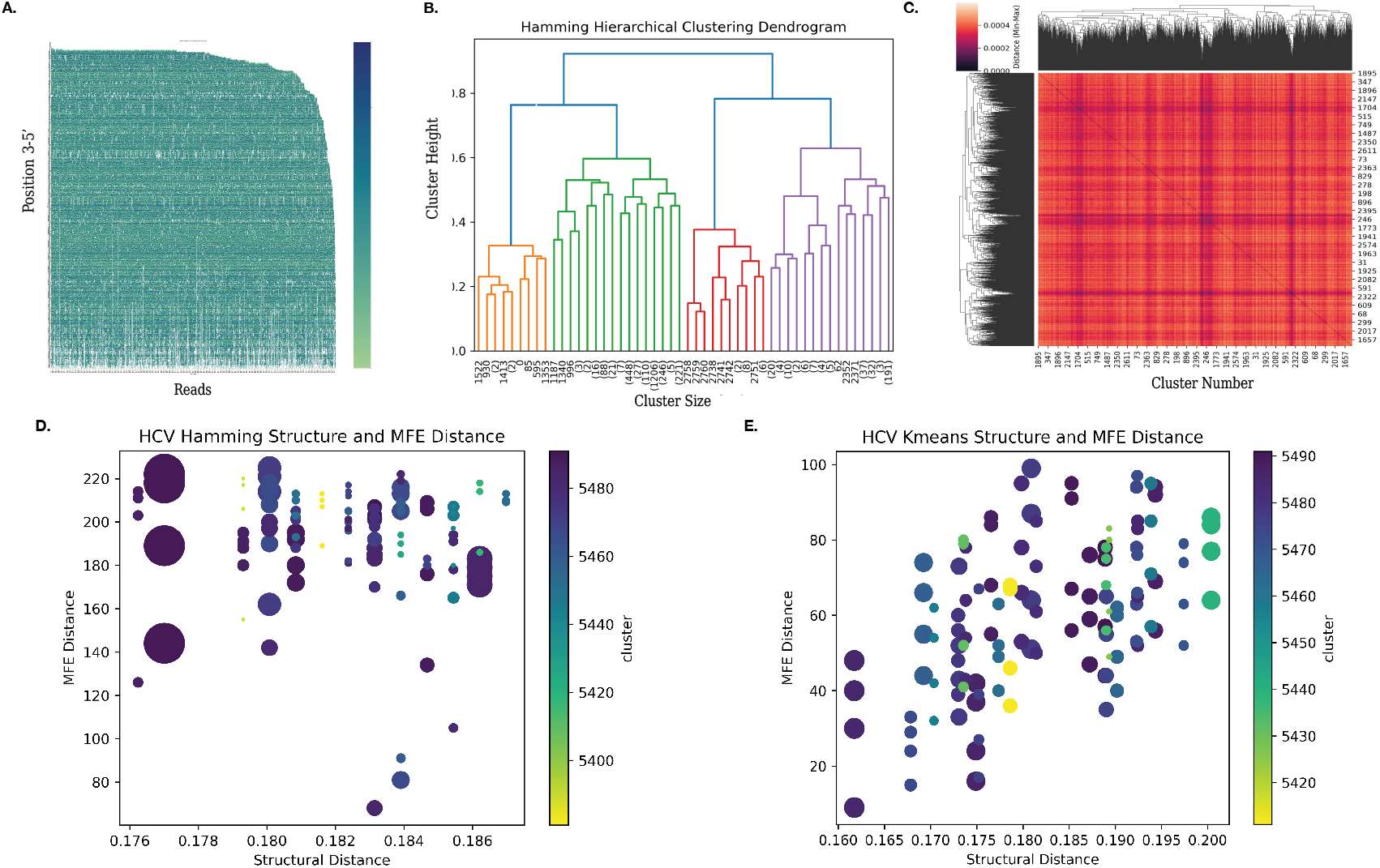
Individual Reactivities and Clustering in Dashing Turtle. Figure shows white areas where 3’ and 5’ regions were shortened or sparse, a noted effect of nanopore sequencing as well as variations in reactivity across the landscape (A). The dendrogram based on the Hamming distance matrix using complete linkage where the maximum distance between observations of pairs of clusters is minimized. The dendrogram is restricted to five levels to capture dominant variations that correspond to the number of alternative structures seen in orthogonal methods (B). HCV IRES cluster mapping for each cluster’s correlation with other clusters in the landscape. Darker areas are less correlated while lighter areas are more strongly correlated. We observe strong correlation with clusters on the right and bottom edge to other clusters in the structural landscape consistent with dominant/native structure (C) Putative structures for HCV IRES and their structural distance from the native HCV IRES in both MFE using Hamming (D) and Kmeans (E) as distance metrics. The size of each point represents the relative cluster size. Each putative structure is represented four times for each of the likely or alternative MFEs as calculated by ViennaRNA. The largest clusters in both methods are structurally close and closest in MFE by a significant degree as expected in the highly structured molecule (D/E)

**Fig. 5.**
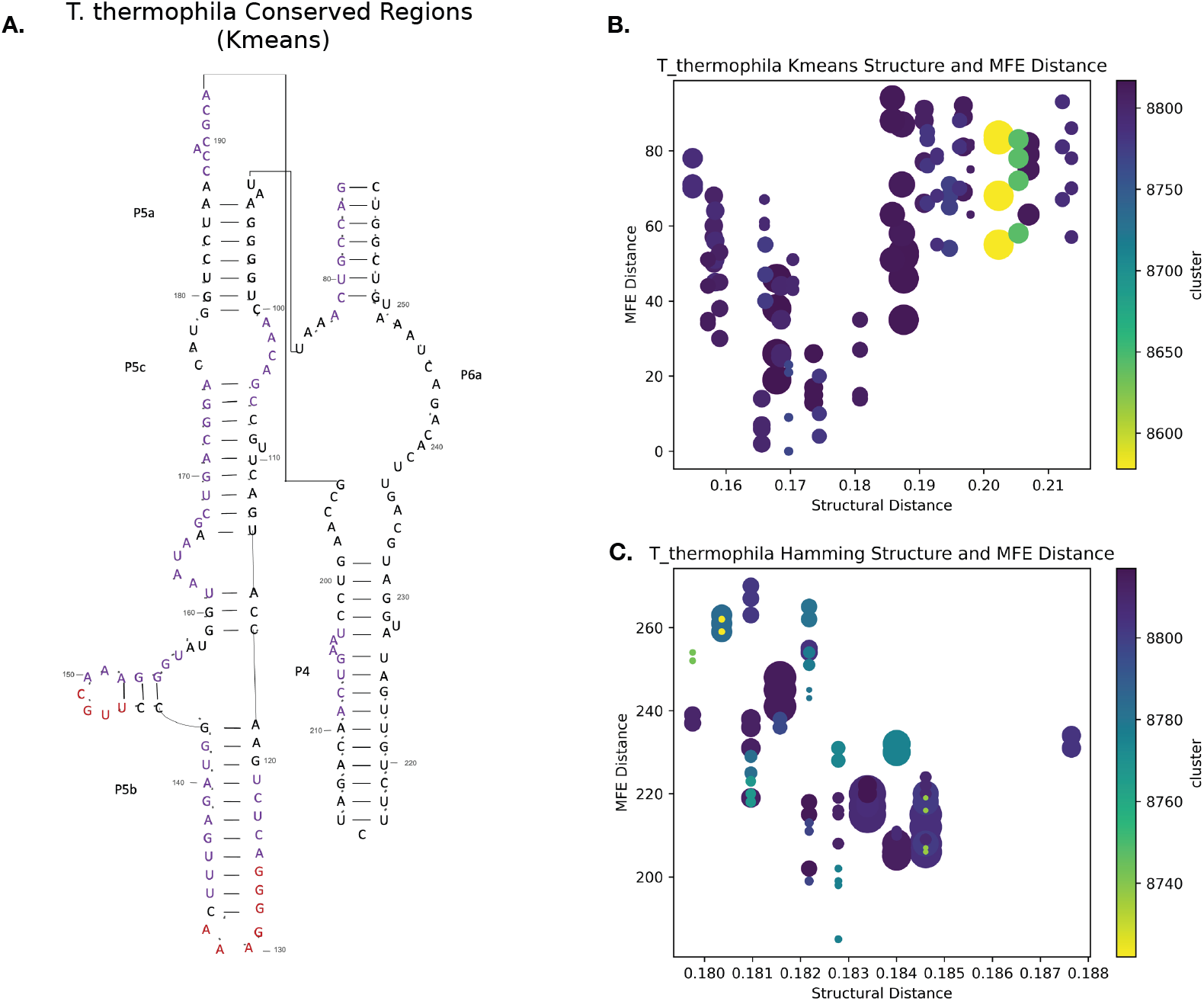
Figure shows conserved regions for kmeans (A). Conserved regions are shown in red (reactive), and purple (non-reactive) for *T. thermophila*. The P4-P6 domain has been observed to fold the fastest and is significantly structurally for catalysis. Here, the Kmeans method aligned most with the P4-P6 region. This was the only region showing significant conservation in the kmeans analysis. Many of the regions indicated to be single stranded are not accessible in the compacted core regions so they are conserved as not reactive (purple). The Hamming based analysis supported this, as there was significant overlap. (B) Putative structures for *T. thermophila* and their structural distance from the native *T. thermophila* psuedoknot free molecule. in both MFE using Kmeans (B) and Hamming (C) and as structural distance metrics. The size of each point represents the relative cluster size. Each putative structure is represented four times for each of the likely or alternative MFEs as calculated by ViennaRNA. The largest clusters are distributed at two structural distances from the native structure. Biologically, *T. thermophila* has been known to forms hinges for splicing. The second dominant cluster suggest a similarly represented alternative conformation (B/C)

#### Clustering

We apply two different distance matrices for clustering: Hamming and Kmeans. Hamming based distance metric applies a threshold to the nucleotide reactivities predicted which filters for high reactivity for each molecule. The distance is the Hamming weight between all reads of the sample (Appendix A.3) and for Kmeans (Appendix A.4), we use standard Euclidean distance. Average read length is calculated for all nanopore reads per sequence, and reads shorter than this read length are removed. We apply agglomerative, hierarchical clustering of the pre-computed distance matrices for both methods. Agglomerative hierarchical clustering is an unsupervised clustering algorithm (Appendix B). Each observation initially forms its own singleton cluster. At each iteration, the two clusters with the smallest dissimilarity are merged, according to our precomputed distance metrics using complete linkage [65, 46]. This process continues iteratively until all observations are grouped into a single cluster, producing a dendrogram that represents the nested relationships among data points. This hierarchical clustering approach parallels the process of RNA folding, in which local secondary structures (such as hairpins and loops) form first, followed by the gradual assembly of larger tertiary structures. Just as agglomerative clustering successively merges small clusters into larger ones, RNA folding proceeds in a hierarchical manner, with local interactions stabilizing initial structures that guide the formation of higher order configurations [66]. The resulting dendrogram of clustering can thus be interpreted as analogous to the folding pathway of the RNA molecule, capturing nested relationships among structural elements. We apply the Elbow method using Within-Cluster-Sum of Squared Errors (WSS) (Appendix B.1) to determine the optimal number of clusters across sequences. While the optimal number varied for each sequence we found a cutoff between 20-30 for each control was effective at highlighting sequence correlations (Figure 5 B./C.) and (Appendix B.2). The averages of these numbers across all sequences was used to determine the cluster number. For all sequences, the optimal average cluster number was 20. We observed that Hamming based clusters have stronger convergence but may obscure subtle variations in the landscape because weaker reactivities near the threshold are pushed to 0. Kmeans based clustering allows investigation of these smaller attributes that are reflected over a gradation. The correlation matrix between clusters help to identify highly anomalous patterns that are less correlated to the rest of the landscape could suggest dis-ordered RNA or intermediate folding, but would require further investigation (Figure 4 C.) [47, 65, 67]. Figure 4 displays the results for HCV IRES.

We calculate the centroids of all clusters using KModes for Hamming based evaluations, where a centroid represents the majority value at a given position within the cluster [68, 69, 70, 71]. The Kmeans centroids correspond to the arithmetic mean of the reads assigned to the individual cluster [47, 67]. The centroid reactivities were scaled to SHAPE reactivities as accepted by ViennaRNA and used in RNAFold to calculate putative structures and the minimum free energies of each putative structure [9]. Table 3 shows the maximum clusters calculated for controls using Kmeans. We observed that highly structured molecules had larger dominant clusters in the landscape, which suggests a preservation of certain structural elements.

**Table 3.** Structural distance reflects the difference between native pseudoknot-free structure and the centroid of the dominant cluster. *δ* MFE reflects the absolute value of the difference between the MFE of the native structure excluding pseudoknots, calculated by ViennaRNA [9]. Synthetic and highly structured RNAs, like riboswitches, have larger dominant clusters than less structured RNAs. *E. coli* tmRNA is the least structured with a 15% dominant cluster, while *T. thermophila*, although highly structured, has two dominant configurations that are almost equal (Figure 5).

| Sequence | Cluster Size | Percent Landscape | Sequence Length | Structural Distance | Native MFE (kcal/mol) | $\vert \Delta \text{MFE}\vert $ Distance (kcal/mol) |
| --- | --- | --- | --- | --- | --- | --- |
| hc16.ligase | 916 | 48.78 | 338 | 79 | -85.50 kcal/mol | 55.10 kcal/mol |
| Lysine riboswitch | 121 | 34.18 | 161 | 47 | -74.60 kcal/mol | 55.58 kcal/mol |
| HCV IRES | 546 | 25.47 | 382 | 119 | -111.50 kcal/mol | 103.35 kcal/mol |
| RNAse_P | 984 | 24.27 | 417 | 106 | -139.90 kcal/mol | 87.25 kcal/mol |
| FMN riboswitch | 19 | 24.05 | 112 | 23 | -21.60 kcal/mol | 9.10 kcal/mol |
| T.thermophila | 356 | 23.94 | 387 | 118 | -104.40 kcal/mol | 79.85 kcal/mol |
| E.coli tmRNA | 379 | 15.03 | 363 | 79 | -105.30 kcal/mol | 81.40 kcal/mol |

### Reconstructing Native Conformations

We compare the MFE of the putative structures with the MFE of the native structure calculated using RNAeval [9]. The MFE distance is the Euclidean distance between the native MFE and the MFE of each of the putative structures. Next, we calculated the structural distance between the native structure and all putative structures using both Hamming weight and Kmeans distance matrices as explained in Section 3.2.2. As Figure 4 D. and E. shows for HCV IRES we found that for all our controls the largest clusters are closest both structurally and in MFE to the native structure. We observed that dominant clusters are those most representative of the native structure. Hamming show larger clusters with fewer variations and better convergence. Kmeans based clusters use the scale of the reactivity and produce much wider variation in the landscape, but predicted MFEs are much closer to the native MFE (Section 3.2.2). In most cases dominant clusters have a minimal structural distance from the native structure and are significantly larger than other patterns in the landscape. Biologically, HCV IRES is highly structured RNA with conserved structural motifs that are functionally necessary for mediating cap-independent translation initiation [72]. Our data exhibits a similarly highly stable dominant conformation (Appendix C.4). Furthermore, we see that most of the structure is represented as conserved regions (discussed in Section 3.4) and are fairly consistent across the landscape. Figure 4 D. shows that for HCV IRES there are several small clusters that are slightly closer in structural distance than the dominant clusters (Appendix C.4). These clusters do not represent significant alternative conformations, but a small change in 1-2 bases from the dominant cluster and could be combined.

We noted that dominant conformations are still less than 50% of the total structural landscape. Jackson et al.’s experimental observations show a similarly that a significant portion of *in vitro* transcripts are subject to misfolding. In particular, *T. thermophila*, a group I self-splicing intron, has a misfolding rate, when transcribed *in vitro*, which correlates with the concentrations of magnesium (Mg +) and other ions. In general, RNA has multiple folding pathways that can result in kinetic partitioning or become trapped in alternative conformations. Our structural landscapes suggest the presence of these molecules in smaller clusters [73].

Luo et al. captured six conformations of *T. thermophila* in cyro-EM experiments in the presence of Mg+ and metal ions, two of these states were in the unlocked positions that had unoccupied active sites, not bound to metal ions. *T. thermophila*’s cluster distribution (Figure 5, also, shows two dominant clusters a short distance (.02) from each other (C.), supporting their observations. Additionally, the dominant structures are slightly further from the native structure than HCV IRES. This is likely because the native structure of *T. thermophila* is usually crystallized in the presence of Mg+.

Overall, the variances observed in our controls from native structures is minimal. Dominant clusters and dominant structures conformed to experimentally confirmed structures and results. Our structural landscapes can represent the native structure as dominant clusters, and could suggest that the other conformations in the landscape represent alternative and/or misfolded conformations. Kmeans based clusters showed the closest MFE to native confirmations for dominant clusters (Table 3).

### Conserved Regions

Conserved regions, are defined here, as three or more contiguous nucleotides where the state is conserved as either reactive or non-reactive for a significant part of the structural landscape. A significantly conserved region is defined as having a prevalence equal to or greater than the dominant structures in the landscape, but the overall putative structure may not be conserved or dominant. For example, significance was set at ≈ 10% of all nucleotides at position *j* in the landscape in Kmeans based clusters and ≈ 40% for Hamming. For HCV IRES because the dominant clusters are ≈ 40% of the structural landscape (Appendix Table 12 and Table 11). Conserved regions tell a story about the accessibility and function of the molecules in the landscape and can identify long range interactions by observing co-occurring conserved regions. These regions may help to identify which regions of the molecule are most critical for preserving function in given states. Specifically, reactive regions imply greater accessibility, while non-reactive regions imply less accessibility. Co-occurring highly reactive regions could help to identify putative binding sites and, if sampled over time, they could provide insight into the hierarchical folding of the molecule.

In our investigation of *T. thermophila*, which has been extensively modeled, it is sequenced without Mg+ and the structure is more accessible than when bound to metal ions and tightly packed. The P4-P6 domains, a 160 nucleotide region of the native structure, have been observed to fold the fastest, and are important to the splicing function. In Figure 5, we detected conserved regions in the P4-P6 domains across the landscape consistent with with experimental analysis. The inaccessible (blue) conserved regions of aligned with experimentally verified packed regions in the tertiary structure. Conserved areas of reactivity (pink) aligned with hinging and locking mechanisms necessary for binding (e.g. magnesium Mg+) and stabilizing the molecule.

The greater variation in *T. thermophila* is correlated with the greater mobility of the molecule necessary for its function, which forms hinges and locks for splicing. Conformations observed during cryo-EM, noted variations near the P1 domain which strongly correlate with our results including the two to three dominant conformations and conserved reactivity [74]. The non-conserved regions cause greater variation in the dominant structures that are not part of the main mechanism within the active site resulting in increased alternative conformations or misfolding [74, 75, 73]. In comparison, the highly structured HCV IRES has conserved regions encompassing almost the entire molecule. In summary, strongly conserved regions correlate well with areas of structural and functional preservation, while low conservation may indicate mis-folded regions or molecules. Our detection of conserved regions parallels both experimental results and correlates with biological function of known interaction mechanisms.

### Discussion & Conclusion

DT is biased for accuracy, but is less sensitive than its comparative algorithms with very low fold change (Figure 2). Comparative algorithms overpredict modifications significantly almost doubling the rate of actual predictions. DT will improve the fold change in the future to maximize coverage.

Ultimately, predicting modifications with ONT technology can be noisy, stochastic process with multiple confounding factors that are not likely to be resolved by conventional approaches. While there has been much emphasis on addressing this issue in recent literature, to date, DT is one of the few that offers a reasonable approach to the challenges of compounded noise, statistical analysis, signal processing and time dependent states with direct sequencing and for prediction of modification on all four nucleotide bases.

DT offers similar performance to existing experimental methods (Section 3.1). We observed accuracy at approximately 50-70% in SHAPE experiments with AcIm, with DT demonstrating the highest average accuracy across our control sequences and comparable performance across all metrics including increased detection of certain structural motifs (Table 2). It is worth noting that accuracy for larger probes on HCV IRES have been reported at 70-90% with 1M7 [8]. We believe an extended investigation in finding optimal probes for chemical modification with ONT technology would be very helpful in increasing the performance across all methods. Additionally, DT offers improved detection of smaller structural motifs such as bulges, and bounds the associated errors rates to its contemporary approaches. Future iterations of DT will be validated on additional chemical probes as they become usable with ONT.

While averaged mutational profiling is useful, it does not provide a comprehensive understanding of the RNA landscape. We have presented clustering methods (Section 3.2.2) using DT that identify dominant conformations in the sample and show that these correlate with known native conformations of these molecules. DT is, also, able to highlight alternative conformations that recapitulate known structural functions (Section 3.3). DT can reasonably reconstruct biological models and could be used to further investigate and model RNA diversity.

Overall, direct RNA sequencing offers possibilities to explore complete transcriptomes and could have significant implications in the understanding RNA dynamics. It is, currently, being used to explore complete transcriptomes and could have significant implications in the understanding RNA dynamics [11, 12, 13]. In the future, it could allow us to more closely observe the dynamic variations of RNA *in vivo*. Structural landscapes may help inform RNA folding mechanisms and offer an understanding of intermediary state changes that could have significant impacts on searches for targets for potential therapeutics.

DT allows for future expansion to include sequence specific biases from ONT data and to more easily account for sequence diversity.

## Data and Source Code Availability

All source code developed for this study is openly available at (DOI: 10.5281/zenodo.XXXXXXX). The repository includes the full implementation of the algorithms, analysis scripts, and example datasets required to reproduce the results presented in this manuscript. The code is released under the MIT License. Additionally, the Dashing Turtle source code is available at and the software package is available at *https://pypi.org/project/DashingTurtle/*

## Author contributions statement

This manuscript and algorithms written and designed by J.W.B, experiments were performed by G.B., and reviewed by E.L. and J.W. The following roles apply: **J.W.B**.: Conceptualization, Data curation, Formal analysis, Methodology, Software, Validation, Visualization, Writing – original draft. **G.B**.: Investigation, Validation, Visualization, Writing – review & editing. **E.L**.: Funding acquisition, Project administration, Resources, Supervision, Writing – review & editing. **J.W**.: Funding acquisition, Project administration, Resources, Supervision, Writing – review & editing.

## Acknowledgments

We thank Dr. Maria Vera Ugalde and Dr. Bruno Sargeuil for generously providing biological resources. We also thank the RNA research community for insightful discussions and feedback.

## Supplementary Data

Supplementary Data are available at NAR Online.

## Definitions, Metrics, and Equations

### Feature Selection and Ablation Analysis

Feature selection plays a critical role in maximizing predictive performance while minimizing overfitting, especially in high-dimensional RNA sequencing datasets. In our study, numerical features were first evaluated using exhaustive feature selection in conjunction with an ensemble learning approach. This was cross-referenced with Fisher’s score to identify the subset of features that most strongly contributed to predicting nucleotide reactivities. Interestingly, the performance difference between the optimal subsets identified by each method was minimal (mean absolute difference ≤ 2%), suggesting that multiple feature combinations can achieve similar predictive power. Both subsets were retained for downstream analyses and were further evaluated using Pearson’s correlation between predicted and experimentally measured reactivities, providing a quantitative metric of model accuracy.

To understand the importance of specific feature groups, we performed an ablation study by selectively removing or isolating individual features. For example, dwell time—a measure of the duration that a nucleotide is recorded in the nanopore—was found to significantly improve predictions when included, as were the basecalling reactivities at each position and the minimum and maximum signals recorded over time. Removing these features individually or in combination resulted in a measurable drop in correlation, highlighting their non-redundant contribution to model performance. This ablation analysis confirms that these features capture critical aspects of nucleotide behavior in nanopore sequencing that are not fully represented by the modification function alone.

Categorical features were evaluated similarly. Although initially included in exhaustive selection, many categorical features introduced bias when the number of sequences was limited. Ablation studies confirmed that removing these features—except for the k-mer attribute—improved model stability without sacrificing predictive accuracy. The k-mer feature was retained due to its robustness with increased observations and its ability to encode local sequence context, which is important for predicting nucleotide interactions.

Overall, the ablation analysis demonstrates that both the choice and weighting of features critically influence predictive performance. Numerical features related to nanopore signal dynamics contribute most directly to model accuracy, while categorical sequence-context features must be carefully curated to avoid introducing bias. These results underscore the importance of systematic feature evaluation and ablation testing to validate that each component meaningfully contributes to model predictions, providing both robustness and interpretability in RNA structural modeling.

### Metric Definitions

The following metrics were used in calculating performance:

- **Accuracy (ACC):** Measures the proportion of correct predictions (both true positives and true negatives) among all predictions.

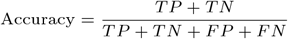
- **Sensitivity (Recall, True Positive Rate, TPR):** Measures the proportion of actual positives correctly identified.

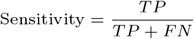
- **F1 Score:** The harmonic mean of precision and sensitivity.

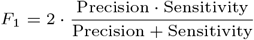
- **Pearson’s Correlation Coefficient (r):** Measures the linear correlation between predicted and true binary labels.

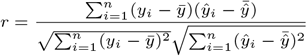
- **Matthew’s Correlation Coefficient (MCC):** A balanced metric considering TP, TN, FP, and FN, informative for imbalanced datasets.

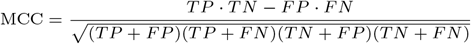

### Hamming Distance

The *Hamming distance* between two sequences of equal length is defined as the number of positions at which the corresponding symbols differ. For sequences *x* = (*x*_1_, *x*_2_, …, *x*_*n*_) and *y* = (*y*_1_, *y*_2_, …, *y*_*n*_), the Hamming distance *d*_*H*_ (*x, y*) is computed as:

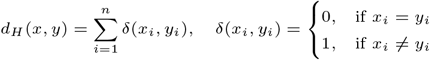

This metric provides a simple measure of sequence dissimilarity suitable for categorical or discrete data.

### K-means Clustering

*K-means clustering* is an unsupervised learning algorithm that partitions a set of *m* observations *{x*_1_, *x*_2_, …, *x*_*m*_*}* into *K* clusters *{C*_1_, *C*_2_, …, *C*_*K*_ *}* such that the within-cluster sum of squares is minimized [76]. Formally, the objective is to find cluster centroids *{µ*_1_, *µ*_2_, …, *µ*_*K*_ *}* that minimize:

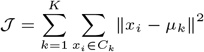

where ∥ ·∥ denotes the Euclidean norm. The algorithm iteratively assigns each observation to the nearest centroid and then updates the centroids as the mean of the assigned points until convergence.

## Agglomerative Hierarchal Clustering

Agglomerative hierarchical clustering is a bottom-up, unsupervised learning algorithm that iteratively merges data points or clusters based on their pairwise similarity until all samples are combined into a single cluster. In contrast to divisive methods, which start from a global cluster and subdivide it, the agglomerative approach begins with each data point as an individual cluster and progressively fuses the most similar pairs.

Formally, given a set of *n* observations *{x*_1_, *x*_2_, …, *x*_*n*_*}*, agglomerative clustering proceeds by:

1. Computing the pairwise distances *d*(*x*_*i*_, *x*_*j*_) between all observations.
2. Merging the pair of clusters (*C*_*p*_, *C*_*q*_) with minimal inter-cluster distance:

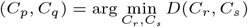

where *D* is the linkage function.

3. Updating the distance matrix and repeating until a single cluster remains.

The linkage criterion determines how distances between clusters are computed, with common options including:

- **Single linkage:** *d*(*A, B*) = min_*a*∈*A,b*∈*B*_ *d*(*a, b*)
- **Complete linkage:** *d*(*A, B*) = max_*a*∈*A,b*∈*B*_ *d*(*a, b*)
- **Average linkage:** 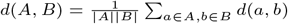
- **Ward’s method:** minimizes the variance within clusters after merging.

### Within-Cluster Sum of Squares (WSS)

The Within-Cluster Sum of Squares (WSS) quantifies the compactness of clusters by measuring the total variance within each group. For a given number of clusters *k*, WSS is defined as

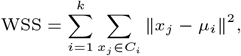

where *C*_*i*_ denotes cluster *i, x*_*j*_ represents individual data points assigned to that cluster, and *µ*_*i*_ is the centroid of *C*_*i*_. Lower WSS values indicate tighter, more homogeneous clusters. WSS was computed across a range of cluster numbers to assess how the internal cohesion of clusters varied with *k*. All clustering and distance computations were performed using the *scikit-learn* library [46].

### Elbow Method

The Elbow method was used to determine the optimal number of clusters based on the WSS metric. As the number of clusters *k* increases, WSS decreases monotonically due to improved cluster fit. However, after a certain point, additional clusters contribute minimal reductions in WSS. The Elbow method identifies this inflection point—often referred to as the “elbow”—where the marginal gain in cluster compactness begins to diminish. This point represents a balance between model complexity and explanatory power. In this study, the elbow was determined by visually inspecting the WSS versus *k* plot to select an appropriate cluster number for downstream structural analyses.

## Tables

### Reagents

**Table 4.**
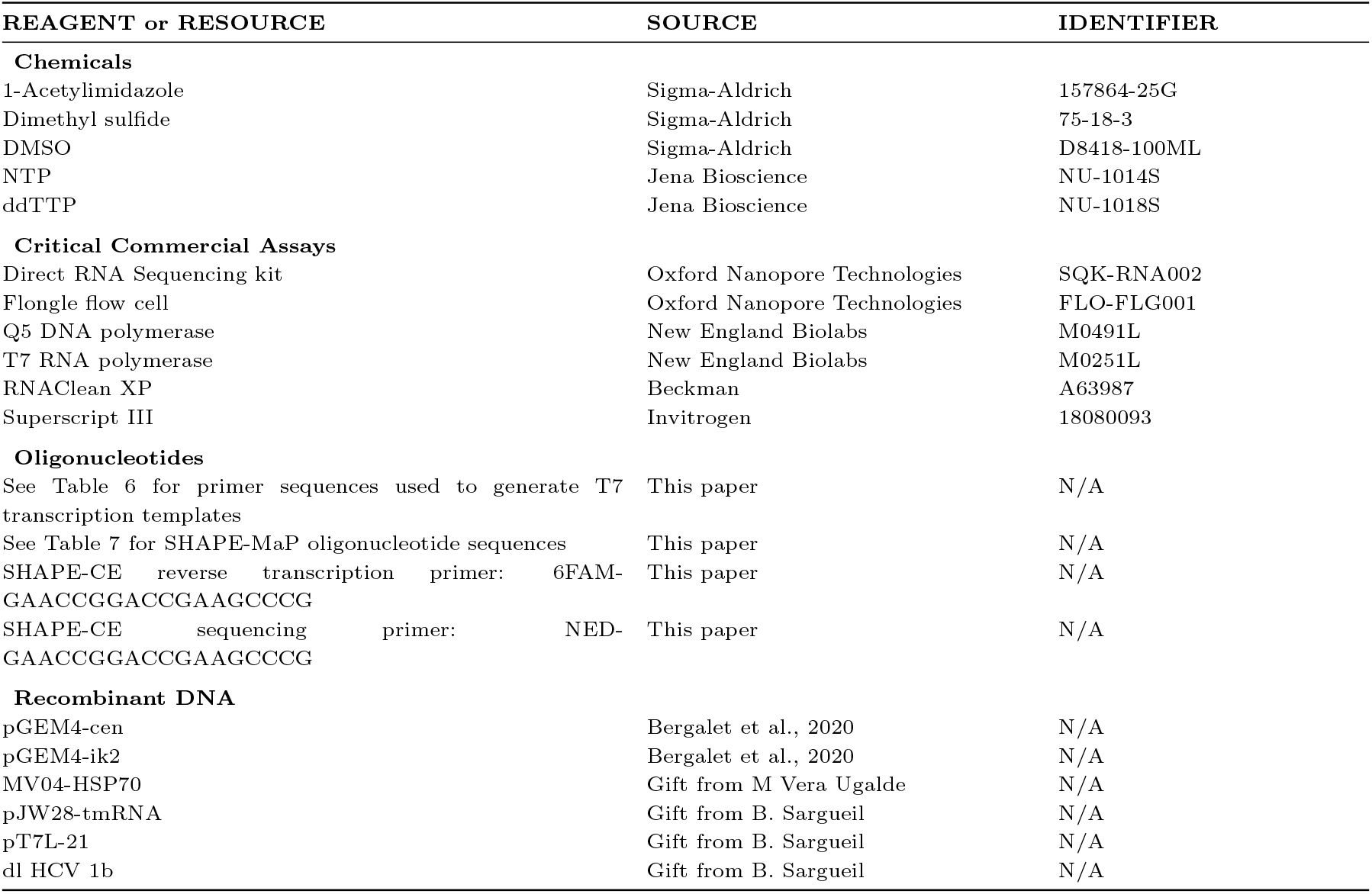
Reagents, commercial assays, oligonucleotides, and recombinant DNA used in this study.

### Oligonucleotides

**Table 6:**
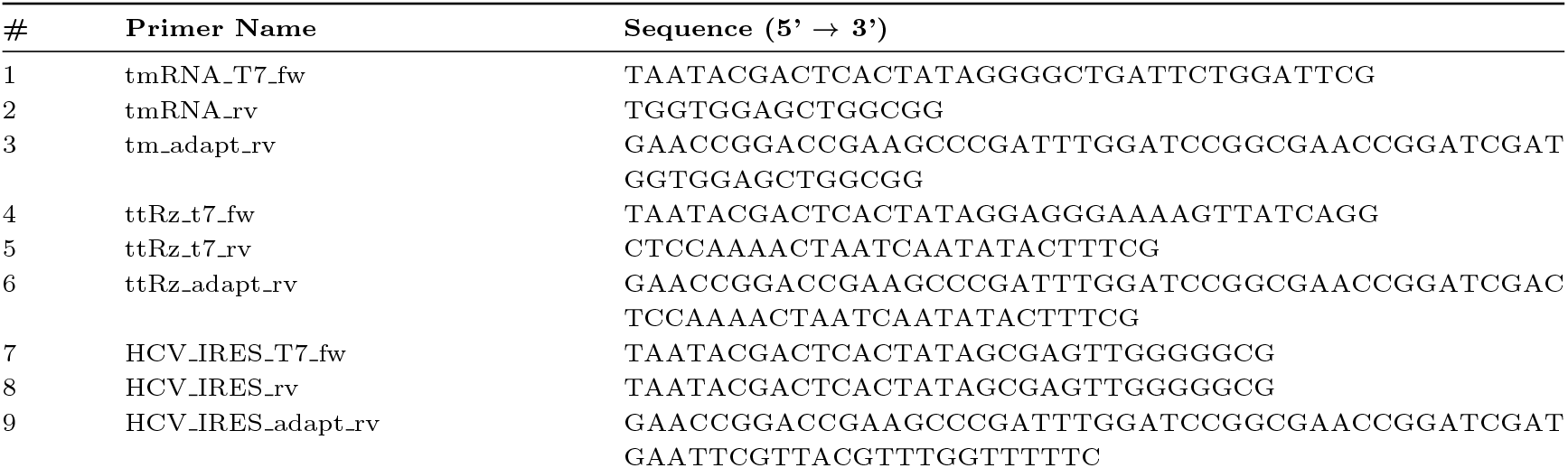
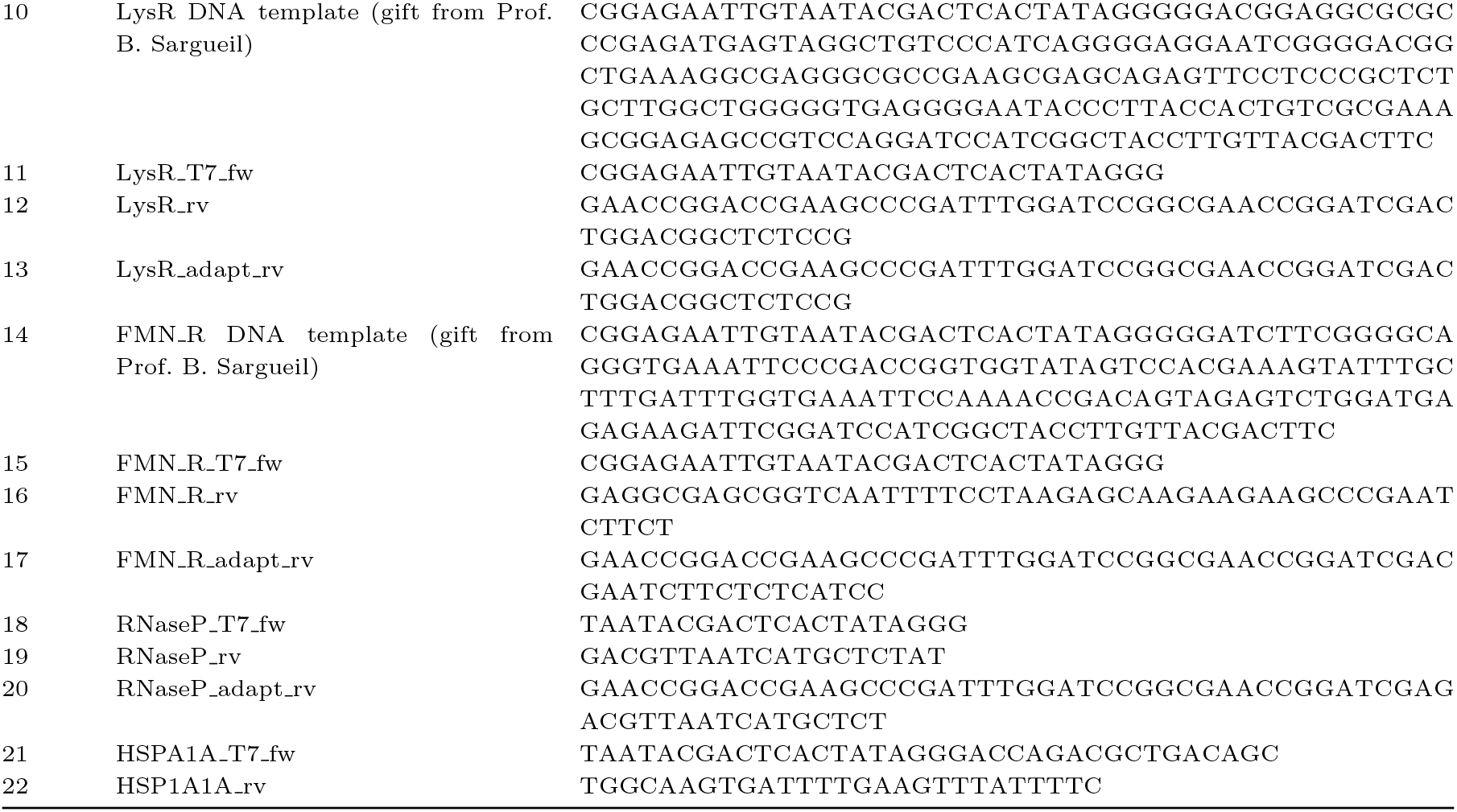
DNA template primers for T7 transcription.

**Table 7:**
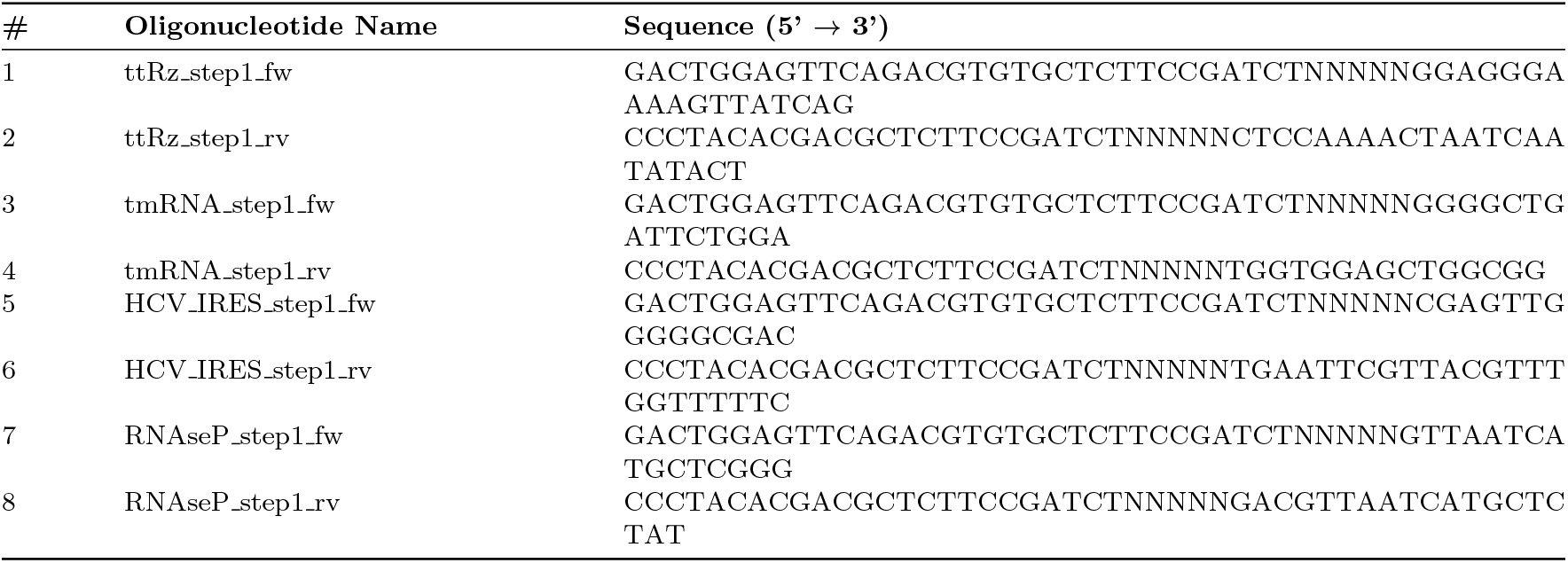
DNA oligonucleotides for SHAPE-MaP.

**Table 8.**
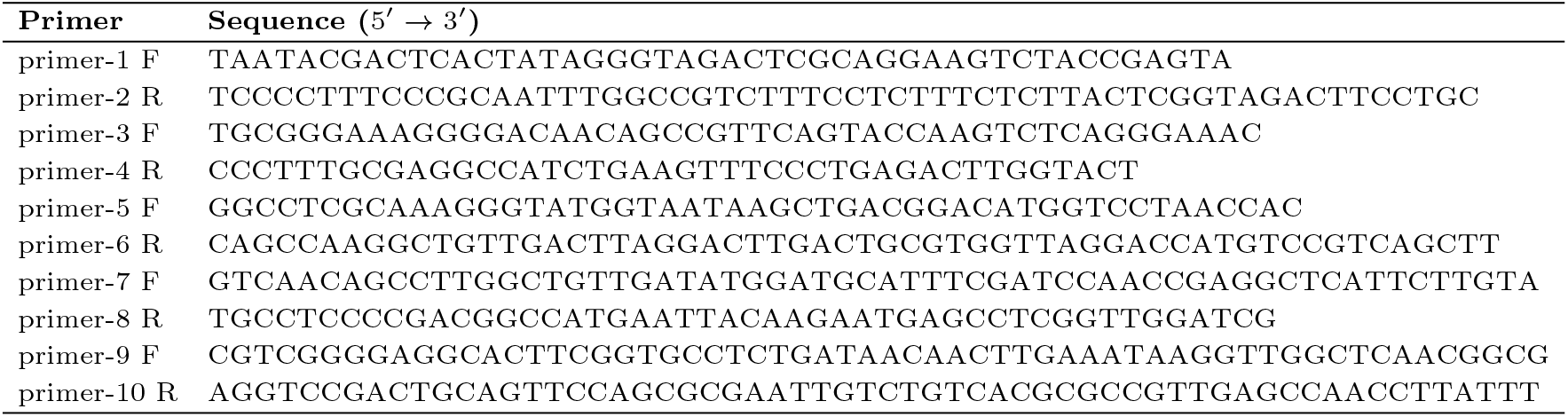
DNA oligonucleotides for hc16 template synthesis.

### Experimental Agreement

**Table 9.** Algorithmic and Experimental Aggreement.

| Predictor | Agree w/<br>SHAPEMAP | Percent Agree<br>SHAPEMAP | Agree w/<br>SHAPECE | Percent Agree<br>SHAPECE | Agree w/<br>Both | Percent Agree<br>Both |
| --- | --- | --- | --- | --- | --- | --- |
| DT | 1717 | **72.26** | 1544 | **67.93** | 1214 | **56.81** |
| RNAcoFold (Using DT) | 1468 | 61.78 | 1356 | 59.66 | 1000 | 46.79 |
| SHAPE-CE | 1537 | 71.92 | 2273 | 100.00 | 1537 | 71.92 |
| PORE-cupine | 1310 | 64.37 | 1157 | 59.49 | 868 | 47.28 |
| nanoSHAPE | 1207 | 50.80 | 1089 | 48.42 | 762 | 35.66 |

Agreement counts and percentages of controls for all algorithms and experiments compared with experiments SHAPEMAP and SHAPECE. Outside of experimental agreement, DT had the highest agreement.

### HCV Clusters

**Table 11:**
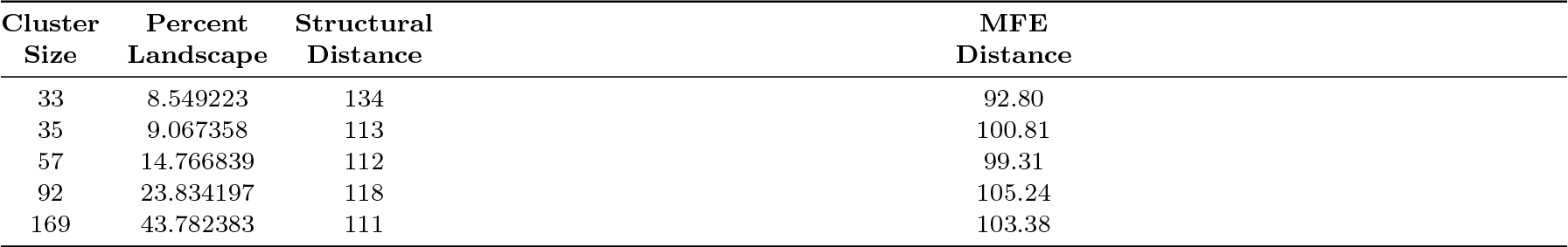
Structural Landscape for HCV IRES (Top 5 Hamming).

| Cluster<br>Size | Percent<br>Landscape | Structural<br>Distance | MFE<br>Distance |
| --- | --- | --- | --- |
| 33 | 8.549223 | 134 | 92.80 |
| 35 | 9.067358 | 113 | 100.81 |
| 57 | 14.766839 | 112 | 99.31 |
| 92 | 23.834197 | 118 | 105.24 |
| 169 | 43.782383 | 111 | 103.38 |

Table 11 shows the top 5 dominant clusters calculated using the Hamming weight of each predicted modification for HCV IRES compared to the native HCV IRES. Structural distance refers to the distance of the putative representative sequence for the cluster from the native sequence. MFE distance calculates the distance of the putative sequence from the native HCV IRES MFE of -111 as predicted by Vienna RNA. Cluster size and percent landscape determine how prevalent the putative sequence is in the landscape. Dominant structures coincide with minimal structural distance from the native structure by a large degree.

**Table 12:**
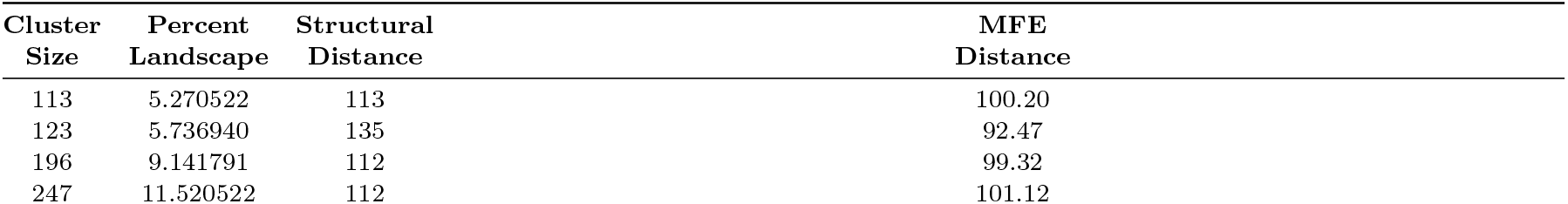
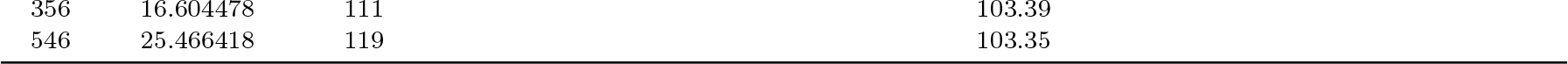
Structural Landscape for HCV IRES (Top 5 kmeans).

Table 12 shows the top 5 dominant clusters found calculated using Kmeans of the reactivity of each predicted modification for HCV IRES compared to the native HCV IRES. Structural distance refers to the distance of the putative representative sequence for the cluster from the native sequence. MFE distance calculates the distance of the putative sequence from the native HCV IRES MFE of -111 as predicted by Vienna RNA. Finally, cluster size and percent landscape determine how prevalent the putative sequence is in the landscape. Dominant structures coincide with minimal structural distance from the native structure by a large degree.

### T.thermophila Clusters

**Table 13:** Structural Landscape for *T. thermophila* (Top 5 Hamming)

| Cluster Size | Percent Landscape | Structural Distance | MFE Distance |
| --- | --- | --- | --- |
| 244 | 9.235428 | 131 | 98.32 |
| 267 | 10.105980 | 136 | 104.08 |
| 327 | 12.376987 | 140 | 102.69 |
| 713 | 26.987131 | 136 | 103.92 |
| 801 | 30.317941 | 136 | 103.07 |

Table 13 shows the top 5 dominant clusters calculated using the Hamming weight of each predicted modification for *T. thermophila* compared to the native *T. thermophila*

**Table 14:** Structural Landscape for *T. thermophila* (Top 5 Hamming)

| Cluster Size | Percent Landscape | Structural Distance | MFE Distance |
| --- | --- | --- | --- |
| 113 | 7.599193 | 140 | 102.40 |
| 119 | 8.002690 | 130 | 104.40 |
| 127 | 8.540686 | 134 | 103.09 |
| 143 | 9.616678 | 134 | 104.38 |
| 356 | 23.940820 | 118 | 79.85 |

Table 14 shows the top 5 dominant clusters calculated using the Kmeans of each predicted modification for *T. thermophila* compared to the native *T. thermophila*

## Figures

### Signal Distribution

**Fig. 6.**
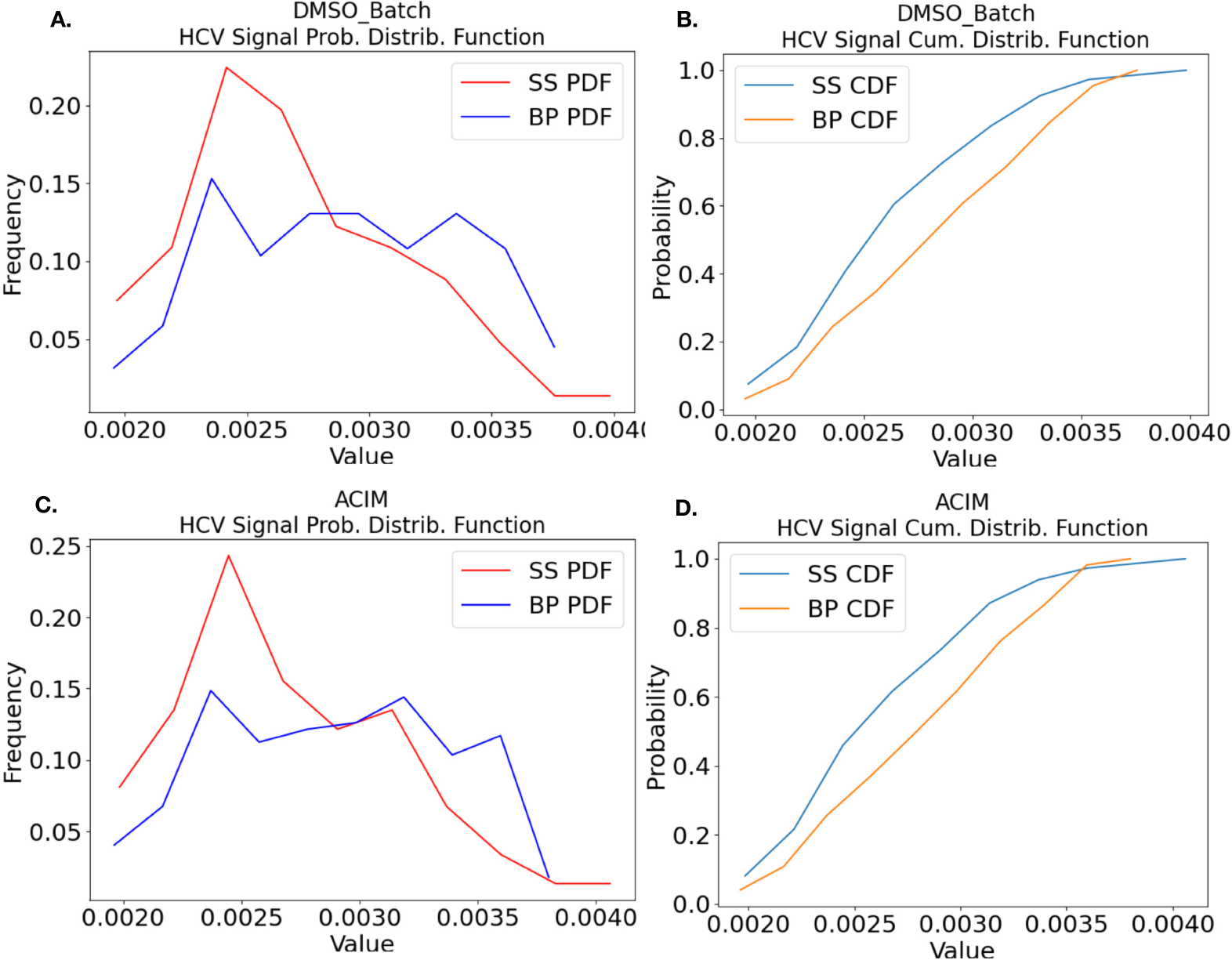
Figure depicts signal distributions for HCV IRES for single stranded (SS) and base paired (BP) nucleotides in native conformation. (A/C) Shows the empirical probability density function for unmodified DMSO and AcIm respectively.(B/D) Shows the empirical cumaltive density function for unmodified DMSO and AcIm respectively. While signal distributions are roughly Gaussian for this sequence they are not unimodal and the distribution will vary based on the sequence context.

### Gaussian Mixture Model (GMM)

Some sequences, were not modified and no changes in cluster size or shape compared to DMSO were observed in those instances or there was significant noise present in the signal. In Figure 7, SHAPE-CE identified a a highly reactive nucleotide position (15), modified by ACIM in a control sequence. In both Kmeans clustering and GMM (Gaussian Mixture Models) a separation into two peaks is observed when ACIM is applied. One peak corresponds roughly to the DMSO peak (7 A./C.) and the second peak represents modification in the input features by modification with ACIM (Figure 7 A./C.).

**Fig. 7.**
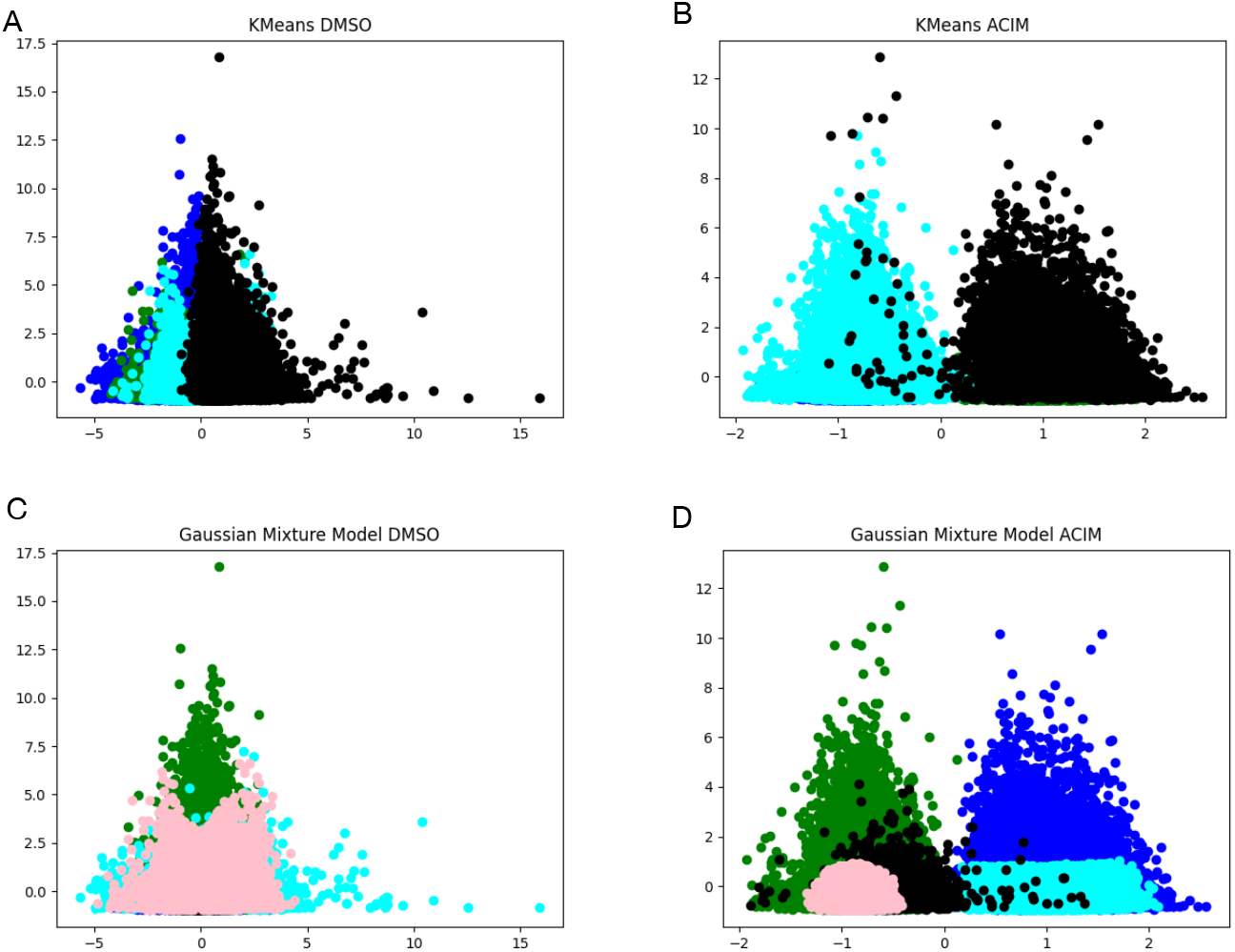
Figure depicts clustering landscape at a modified position. A) Shows Kmeans clustering of unmodified position and corresponds roughly to a Guassian distribution we see this also in the histogram of unmodified sequences. B) Shows KMeans clustering under modification with ACIM two distinct peaks are clearly visible. C./ D. Shows clustering using a Gaussian Mixture Model to decompose and deconvolute the clusters. Here we see 3 dominant clusters at the nucleotide position.

Figure 7 D, we also observed an additional cluster (cyan) in the modified peak (blue), possibly indicating alternative structures of cen in solution when modified. For instance, some strands may have been intra-moleculary paired at position 15 reducing the reactivity at that position; however, they were still modified by ACIM. When clusters are visualized over the entire sequence we are able to identify several possible alternative configurations in solution. The most dominant, or highest frequency of occurrence would form dominant modification peaks as we see in Figure 7 B and D. and within those peaks we can observe additional clustering that may correspond to alternate conformations (D.).

### Conserved Regions

**Fig. 8.**
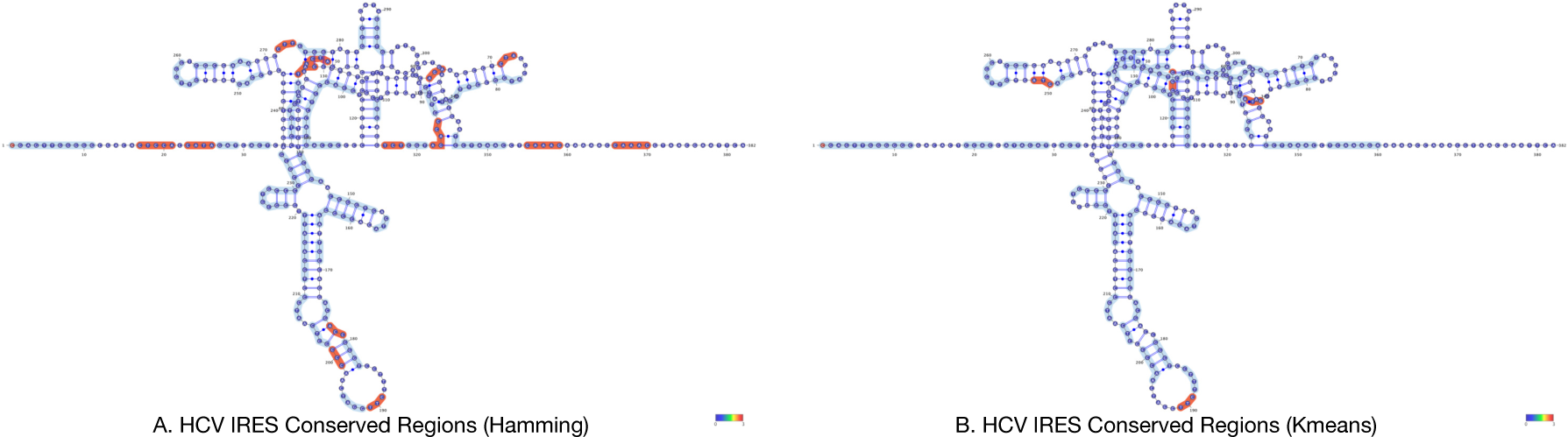
Figure shows conserved regions for both hamming (A) and kmeans (B). Reactive regions are in pink, and non-reactive regions are in blue for HCV IRES. HCV IRES is highly structured RNA and most of the bases are can be represented as conserved regions. Structural motifs are highly conserved as they are functionally necessary for mediating mediates cap-independent translation initiation [72]

